# Topographical and cell type-specific connectivity of rostral and caudal forelimb corticospinal neuron populations

**DOI:** 10.1101/2023.11.17.567623

**Authors:** Lina Marcela Carmona, ET Thomas, Kimberly Smith, Bosiljka Tasic, Rui M. Costa, Anders Nelson

## Abstract

Corticospinal neurons (CSNs) synapse directly on spinal neurons, a diverse group of neurons with unique structural and functional properties necessary for body movements. CSNs modulating forelimb behavior fractionate into caudal forelimb area (CFA) and rostral forelimb area (RFA) motor cortical populations. Despite their prominence, no studies have mapped the diversity of spinal cell types targeted by CSNs, let alone compare CFA and RFA populations. Here we use anatomical and RNA-sequencing methods to show that CSNs synapse onto a remarkably selective group of spinal cell types, favoring inhibitory populations that regulate motoneuron activity and gate sensory feedback. CFA and RFA CSNs target similar spinal cell types, with notable exceptions that suggest these populations differ in how they influence behavior. Finally, axon collaterals of CFA and RFA CSNs target similar brain regions yet receive surprisingly divergent inputs. These results detail the rules of CSN connectivity throughout the brain and spinal cord for two regions critical for forelimb behavior.

## INTRODUCTION

Voluntary movement emerges from activity widely distributed throughout the nervous system^1^. Signals generated by the brain recruit neuronal circuits in the spinal cord that control body muscles and regulate sensory feedback^2^. These spinal circuits are derived from six major developmentally defined classes of dorsal neurons and five classes of ventral neurons, each of which comprise several transcriptionally defined subtypes with distinct structural and physiological properties^3–5^. The brain shapes body movement in large part through the connections made with this diverse pool of spinal neurons, enabling countless patterns of muscle activation and regulating the structure of sensory feedback. Among the neuronal populations that project to the spinal cord, corticospinal neurons (CSNs) play an important role in this transformation^6^.

CSNs are the most populous descending input to the spinal cord^7^. The primary axons of CSNs are widely distributed, forming terminal fields that collectively innervate nearly the entire rostro-caudal extent of the spinal cord^8–10^. CSN axons also span the dorso-ventral extent of the spinal grey, allowing them to synapse on diverse neuron subtypes distributed across spinal laminae. Indeed, targeted anatomical tracing efforts using transsynaptic tracing or appositional analysis reveal several spinal neuron subtypes that receive input from CSNs, including premotor interneurons and sensory relay neurons^7,11,12^. And yet, spinal neurons are incredibly diverse in identity, posing a challenge for comprehensively mapping the full matrix of corticospinal connectivity. Moreover, CSNs can be divided based on their location in sensorimotor cortex, and the rules of spinal connectivity may differ across these populations^7^. Specifically, the cell bodies of CSNs that are involved in forelimb control are distributed across two regions of motor cortex: caudal forelimb area (CFA) and rostral forelimb area (RFA)^13^. Both CFA and RFA are involved in controlling arm and paw movements in rodents, but how their connectivity with spinal neuron subtypes is defined remains unknown^8,14,15^. CSNs also produce robust axon collaterals that innervate many supraspinal regions, an architecture that enables coordination of behavioral signals across the entire nervous system^7,16^. How CSNs in CFA and RFA differ in their collateral connectivity with these brain regions has also been incompletely described^17^. Finally, the roles of neurons are defined in part by their synaptic inputs, and we know little about the similarities and differences in brainwide input to CFA and RFA CSNs.

Resolving these principles of connectivity is essential to understand how descending motor control pathways enable behavior. In this study, we used a suite of transneuronal tracing, single-nucleus RNA-sequencing, and synaptic electrophysiology to reveal the topographical and cell-type connectivity of RFA and CFA populations. We characterized the innervation strategies of CFA and RFA CSNs in the spinal cord, revealing the cellular networks that ultimately shape descending behavioral commands. We further mapped both the brainwide axon collateral and presynaptic input structures of CFA and RFA CSNs, discovering surprising differences in their organization that suggest unique functional roles in behavior. Together, these results reveal the infrastructure of two unique parallel descending motor control pathways.

## RESULTS

### CFA and RFA corticospinal neurons target neurons in distinct spinal regions

Corticospinal neurons (CSNs) are widely distributed across the cortical mantle, including across distinct sensorimotor cortical regions^7,18^. Two major populations of motor CSNs arise from the caudal forelimb area (CFA) and rostral forelimb area (RFA) of motor cortex^13,19^. We confirmed this by injecting a retrogradely-transported adeno-associated virus (AAV) encoding a fluorescent reporter (AAV-retro-FP) into segments C3-C8 of the cervical spinal cord, home to the circuits that regulate forelimb control (Figure 1A, N = 3)^20^. We then imaged antibody-enhanced fluorophore labeling and used the imaging analysis pipeline BrainJ to map their positions to a common brain atlas^21^. Visual inspection of the dorsal surface of the brain revealed two physically separated groups of CSNs with positions corresponding to CFA and RFA (**Figure 1B**)^19^. CFA an RFA CSNs project to overlapping, but partially distinct regions of the spinal cord^8,9^. We confirmed this using an intersectional strategy to express different fluorophores exclusively in CFA and RFA CSNs (**Figure 1C**, N = 6, from same mice as in Figure 3). We first injected a 1:1 cocktail of AAV-retro-Cre and AAV-retro-FlpO in the cervical spinal cord. We followed this with injections of AAV-FLEX-GFP into CFA and AAV-FRT-tdTomato into RFA, labeling CFA and RFA axonal projections with GFP and RFP, respectively (**Figure 1C**). Visual inspection of the cervical spinal cord revealed unique projection patterns of CFA and RFA CSNs. RFA CSNs appeared to target deeper regions of the spinal cord, and CFA CSNs more superficial regions, consistent with previous studies (**Figure 1D**)^8,9^.

**Figure 1.**
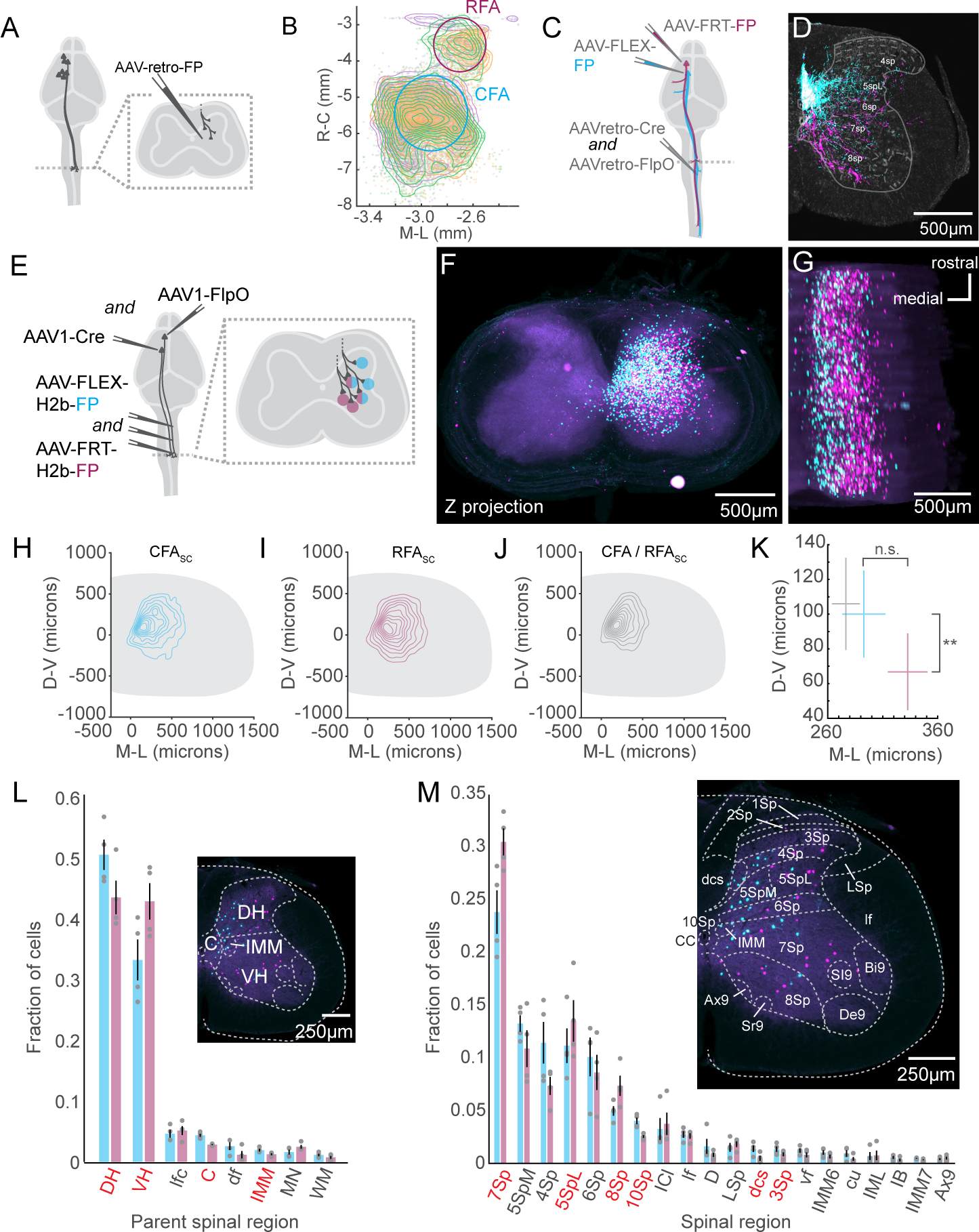
CFA and RFA corticospinal neurons target neurons in distinct spinal regions. **A)** Strategy to label brainwide inputs to the cervical spinal cord. **B)** Top-down view of a 3D reconstruction of the mouse brain showing CSNs as individual points. A contour map of CSN cell bodies is superimposed. The approximate locations of CFA and RFA are indicated. The three colors indicate samples from 3 separate experiments. **C)** Strategy to simultaneously label CFA and RFA CSNs. **D)** Photomicrograph of a transverse section of the spinal cord, with fluorescently labeled CFA (cyan) and RFA (magenta) axons. Neurotrace is in grey. Major spinal laminae are demarcated (representative of N = 6). **E)** Strategy to simultaneously label spinal neurons that are innervated by CFA or RFA. **F)** Z stack projection of aligned photomicrographs of the cervical spinal cord. CFA_SC_ is in cyan, RFASC is in magenta. **G)** Rotated 3D reconstruction of a section of the cervical spinal cord. **H)** Contour plot of CFA_SC_ labeling. D-V is dorsoventral; M-L is mediolateral. **I)** Contour plot of RFA_SC_ labeling. **J)** Contour plot of double labeled neurons (CFA/RFA_SC_ neurons). **K)** Mean centroid positions of CFA_SC_, RFA_SC,_ CFA/RFA_SC_ populations. Errorbars depict SEM. **L)** The major spinal cord structures containing CFA_SC_ and RFA_SC_ neurons. Regions colored red indicate a significant difference between CFA_SC_ and RFA_SC_ fractions. The inset is a photomicrograph with major spinal structures indicated. **M)** The spinal cord laminae and nuclei containing CFA_SC_ and RFA_SC_ neurons. The inset is a photomicrograph with spinal structures indicated. Refer to Table 1 for a complete list of the spinal cord structures and their corresponding acronyms.

**Table 1.**
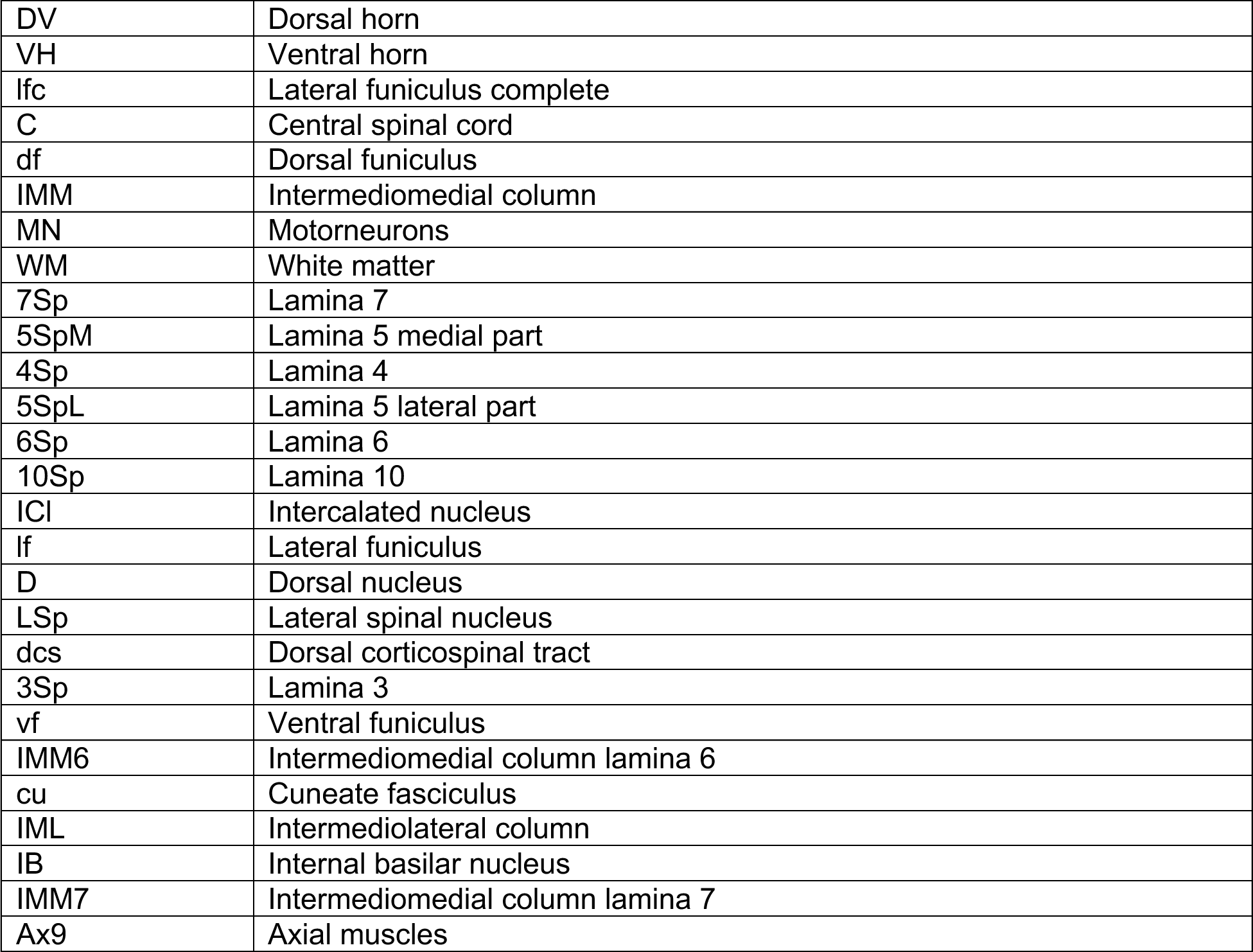

|  |  |
| --- | --- |
| DV | Dorsal horn |
| VH | Ventral horn |
| lfc | Lateral funiculus complete |
| C | Central spinal cord |
| df | Dorsal funiculus |
| IMM | Intermediomedial column |
| MN | Motorneurons |
| WM | White matter |
| 7Sp | Lamina 7 |
| 5SpM | Lamina 5 medial part |
| 4Sp | Lamina 4 |
| 5SpL | Lamina 5 lateral part |
| 6Sp | Lamina 6 |
| 10Sp | Lamina 10 |
| ICI | Intercalated nucleus |
| lf | Lateral funiculus |
| D | Dorsal nucleus |
| LSp | Lateral spinal nucleus |
| dcs | Dorsal corticospinal tract |
| 3Sp | Lamina 3 |
| vf | Ventral funiculus |
| IMM6 | Intermediomedial column lamina 6 |
| cu | Cuneate fasciculus |
| IML | Intermediolateral column |
| IB | Internal basilar nucleus |
| IMM7 | Intermediomedial column lamina 7 |
| Ax9 | Axial muscles |

The presence of CFA and RFA CSN axons in distinct laminae of the spinal cord may reflect the innervation of topographically distinct neurons or may reflect differences in positional innervation of somatodendritic arbors. We disambiguated these possibilities by labeling the spinal neurons targeted by CFA and RFA CSNs, using anterograde transneuronal tracing^22,23^. First, we injected a cocktail of AAV-FLEX-H2b-GFP and AAV-FLEX-H2b-RFP throughout the cervical spinal cord followed by injections of high titer AAV1-Cre and AAV1-FlpO into CFA and RFA, respectively (**Figure 1E**, N = 4). This led to the transneuronal spread of Cre and FlpO recombinases, and the expression of GFP and RFP in spinal somata targeted by CFA and RFA, respectively. We imaged antibody-enhanced fluorophore labeling and leveraged the image reconstruction and analysis pipeline SpinalJ to register transverse cervical spinal cord segments^24^. This revealed a volume of cervical tissue with large populations of CFA_SC_ and RFA_SC_ neurons (**Figure 1F-G**, CFA_SC_: 1052±223.96 cells, RFA_SC_1568±579.12 cells). Measuring the positions of CFA_SC_ and RFA_SC_ neurons revealed a great deal of overlap between these populations, including a fraction of neurons innervated by both CFA and RFA (**Figure 1H-J**, 75±32.12 double-labeled cells). However, mapping the centroids of CFA_SC_ and RFA_SC_ populations across animals revealed a significant difference in the positions of innervated neurons: RFA CSNs target more ventral regions of the spinal grey, while CFA CSNs target superficial regions (**Figure 1K**, Coordinates relative to the central canal. CFA_SC_: 292.74±19.70µm M/L, 100.03±25.11µm D/V; RFA_SC_: 332.75±18.15µm M/L, 66.74±22.18µm D/V: D/V: p = 0.0081, paired t-test, M/L: p = 0.1504, paired t-test). Interestingly, double-labeled neurons settle in medial regions of spinal cord, indicating convergent input is constrained to a select region of the spinal cord (CFA/RFA_SC_: 276.40±12.70µm M/L, 105.91±26.51µm D/V).

The spatial coordinates of the spinal cord map onto well-defined and functionally distinct spinal laminae and nuclei^25^. For example, the dorsal horn is home to several substructures involved in shaping sensory feedback, while the ventral horn is home to motor pools and the premotor neurons involved in shaping muscle activity, amongst other functions^20^. With this knowledge, we again used SpinalJ to map the positions of GFP^+^ and RFP^+^ spinal neurons to a common spinal cord atlas. We discovered a greater fraction of RFA_SC_ neurons in the ventral horn compared to CFA_SC_ neurons (CFA_SC_: 0.331±0.033, RFA_SC_: 0.427±0.030, p = 0.0004, paired t-test). Conversely, a greater fraction of CFA_SC_ neurons is found in the dorsal horn (CFA_SC_: 0.503±0.025, RFA_SC_: 0.434±0.028, p = 0.0025, paired t-test) (**Figure 1L**). We next performed a granular analysis, identifying individual laminae home to spinal neurons targeted by CFA or RFA. Among spinal regions with labeled neurons, lamina 7, lateral lamina 5, and lamina 8 are home to a greater fraction of RFA_SC_ neurons compared to CFA_SC_ neurons (CFA_SC_ versus RFA_SC_: 7Sp: 0.239±0.021 v. 0.306±0.013, p = 0.007; 5SpL: 0.112±0.016 v. 0.135±0.020, p = 0.027; 8Sp: 0.050±0.005 v. 0.074±0.010, p = 0.021, paired t-tests), while lamina 10, and to a lesser extent lamina 3 contain a greater fraction of CFA_SC_ neurons (**Figure 1M**, 10Sp: 0.041±0.003 v. 0.026±0.001, p = 0.008; dcs: 0.014±0.003 v. 0.005±0.002, p = 0.041; 3Sp: 0.014±0.003 v. 0.009±0.002, p = 0.024, paired t-tests). These results establish the topography of corticospinal connectivity and discover that CFA and RFA CSNs target overlapping but distinct regions of the spinal cord. Dorsal horn neurons are more likely to receive CFA input and ventral horn neurons more likely to receive RFA input, with individual spinal laminae contributing to this topographical bias.

### CSNs are selective in their spinal neuron targets

The topography of spinal neurons reflects their transcriptional identity, electrophysiological properties, and synaptic connectivity. For instance, many spinal populations are defined by the expression of 2-3 transcription factors, settle in compact regions of the spinal cord, and are selective in their connectivity with sensory-motor circuits^5,26,27^. The molecular and functional diversity of these spinal populations enables flexible control of motor output and precision processing of sensory feedback^28^. What spinal subtypes do corticospinal neurons target? Previous efforts relied on assessing connectivity between CSNs and individual neuronal subtypes^11,12^, an approach that is difficult to scale to a more complete representation of spinal cell type diversity. Instead, we took a higher throughput approach, AnteroT-seq, by combining anterograde transneuronal cellular labeling with single-nucleus RNA sequencing. We injected high-titer AAV1-Cre into CFA or RFA of CAG-Sun1-sfGFP (INTACT) mice^29^, which express nuclear-localized GFP in the presence of Cre recombinase. This approach results in the transneuronal expression of Cre, thus GFP, in spinal neurons targeted by CFA or RFA (**Figure 2A**). We first confirmed that this approach labeled spinal neurons in a fashion consistent with the results from Figure 1. Visual inspection of antibody-enhanced INTACT labeling in spinal sections (**Figure 2B**) and volumetric reconstructions (**Figure 2C**) confirmed GFP expression throughout the spinal grey. We quantified the distribution of GFP-labeled nuclei and confirmed that RFA_SC_ nuclei are more ventrally positioned compared to CFA_SC_ nuclei, like what we observed in our earlier efforts (**Figure 2D**, CFA: 100.67±5.89μm D/V, RFA: 60.03±5.93μm D/V, p = 1.2 × 10^−6^, unpaired t-test). Next, fresh-frozen cervical spinal cords were dissociated and nuclei with GFP and DAPI fluorescence were enriched using fluorescence activated cell sorting (FACS). Individual nuclei were profiled using SMART-Seq v4 yielding a total of 765 and 678 nuclei post QC for CFA and RFA injected samples, respectively. We achieved a similar median gene detection for each target group with 4769 genes for the CFA population and 4415 genes for the RFA population. Louvain clustering of the aggregate data identified 9 clusters, which, as expected, consisted primarily of neurons (**Figure S1A,B**).

**Figure 2.**
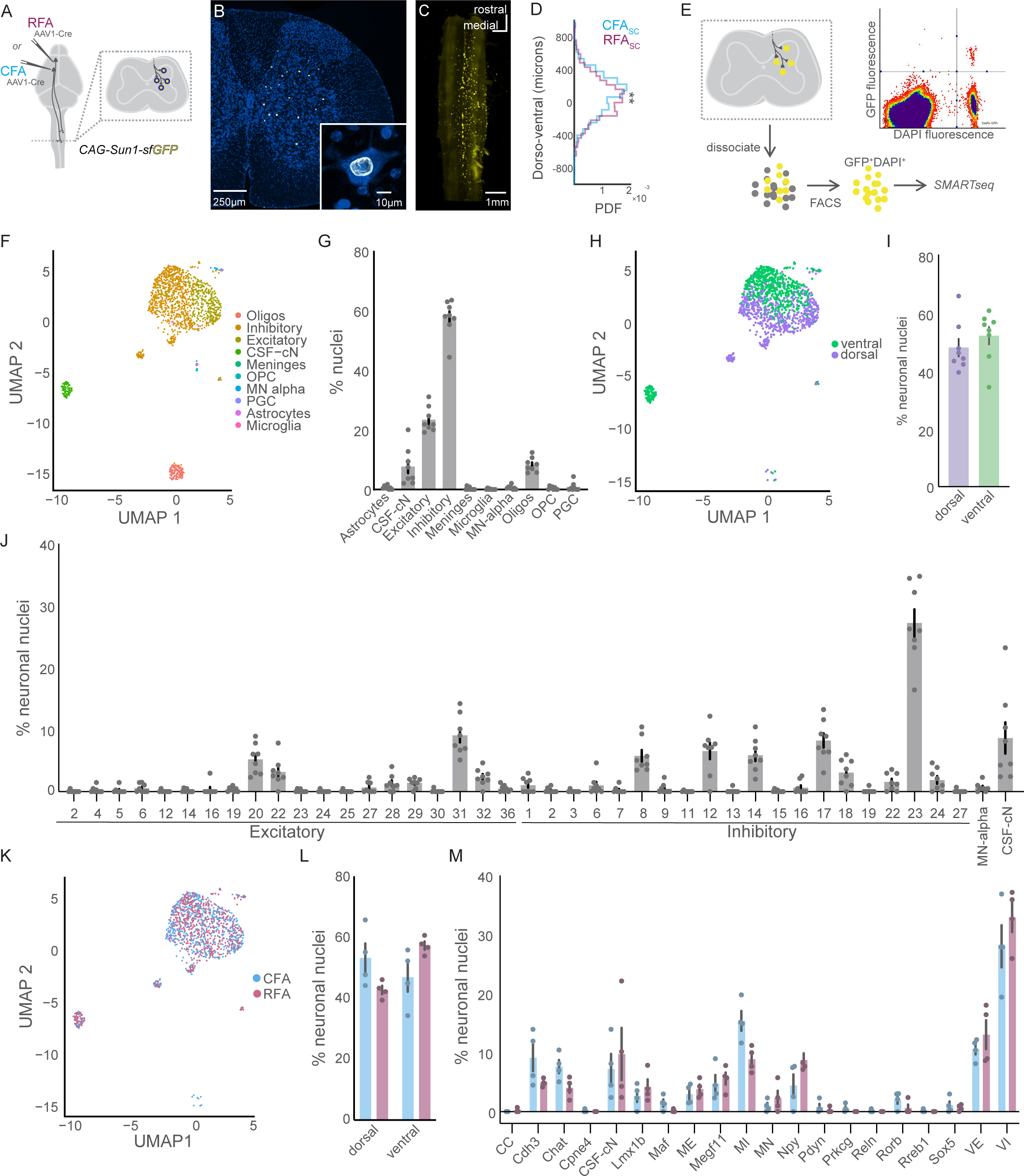
CSNs are selective in their spinal neuron targets. **A)** Strategy to label spinal cord nuclei targeted by CFA or RFA with GFP. **B)** Exemplar photomicrograph of CFA_SC_ nuclei in the spinal cord. **C)** Light sheet microscopy reconstruction of CFA_SC_ nuclear labeling. **D)** Histogram of the dorso-ventral distribution of CFA_SC_ and RFA_SC_ labeling. **E)** Strategy to isolate and perform RNA-seq on CFA_SC_ or RFA_SC_ neurons. An exemplar FACS plot is shown, depicting GFP and DAPI co-labeled nuclei in the top right quadrant. **F)** UMAP plot of all CSN_SC_ sequenced nuclei passing quality control mapped to the major cell types of the harmonized spinal cord atlas from Russ et al. **G)** The percent of nuclei belonging to each major cell type, across individual experiments. **H)** UMAP plot with nuclei labeled by dorsal and ventral positional identity. **I)** The positional identity of nuclei across individual experiments. **J)** The percent of neuronal nuclei belonging to harmonized spinal neuron type clusters, across individual experiments. **K)** UMAP plot of sequenced nuclei labeled by CFA or RFA experimental condition. **L)** The positional identity of CFA_SC_ and RFA_SC_ neurons, across individual experiments. **M)** The percent of neuronal nuclei belonging to the harmonized spinal neuron families across individual experiments. In all plots, error bars indicate SEM. Refer to Table 2 for a complete list spinal cell types, families, and their corresponding acronyms.

**Table 2.**

|  |  |
| --- | --- |
| CSF-cN | Cerebrospinal fluid contacting neurons |
| MN-alpha | Motor neurons – alpha |
| Oligos | Oligodendrocytes |
| OPC | Oligodendrocyte precursor cells |
| PGC | Preganglionic cells |
| CC | Clarke's Column |
| Cdh3 | Cad3 dorsal inhibitory |
| Chat | Chat |
| Cpne4 | Cpne4 dorsal excitatory |
| Lmx1b | Lmx1b mid excitatory |
| Maf | Maf dorsal excitatory |
| ME | Mid excitatory |
| Megf11 | Megf11 dorsal and mid excitatory |
| MI | Mid inhibitory |
| MN | Motor neurons |
| Npy | Npy dorsal inhibitory |
| Pdyn | Pdyn dorsal inhibitory |
| Prkcg | Prkcg dorsal excitatory |
| Reln | Reln dorsal excitatory |
| Rorb | Rorb dorsal inhibitory |
| Rreb1 | Rreb1 dorsal excitatory |
| Sox5 | Sox5 dorsal excitatory |
| VE | Ventral excitatory |
| VI | Ventral inhibitory |

**Table 3.**
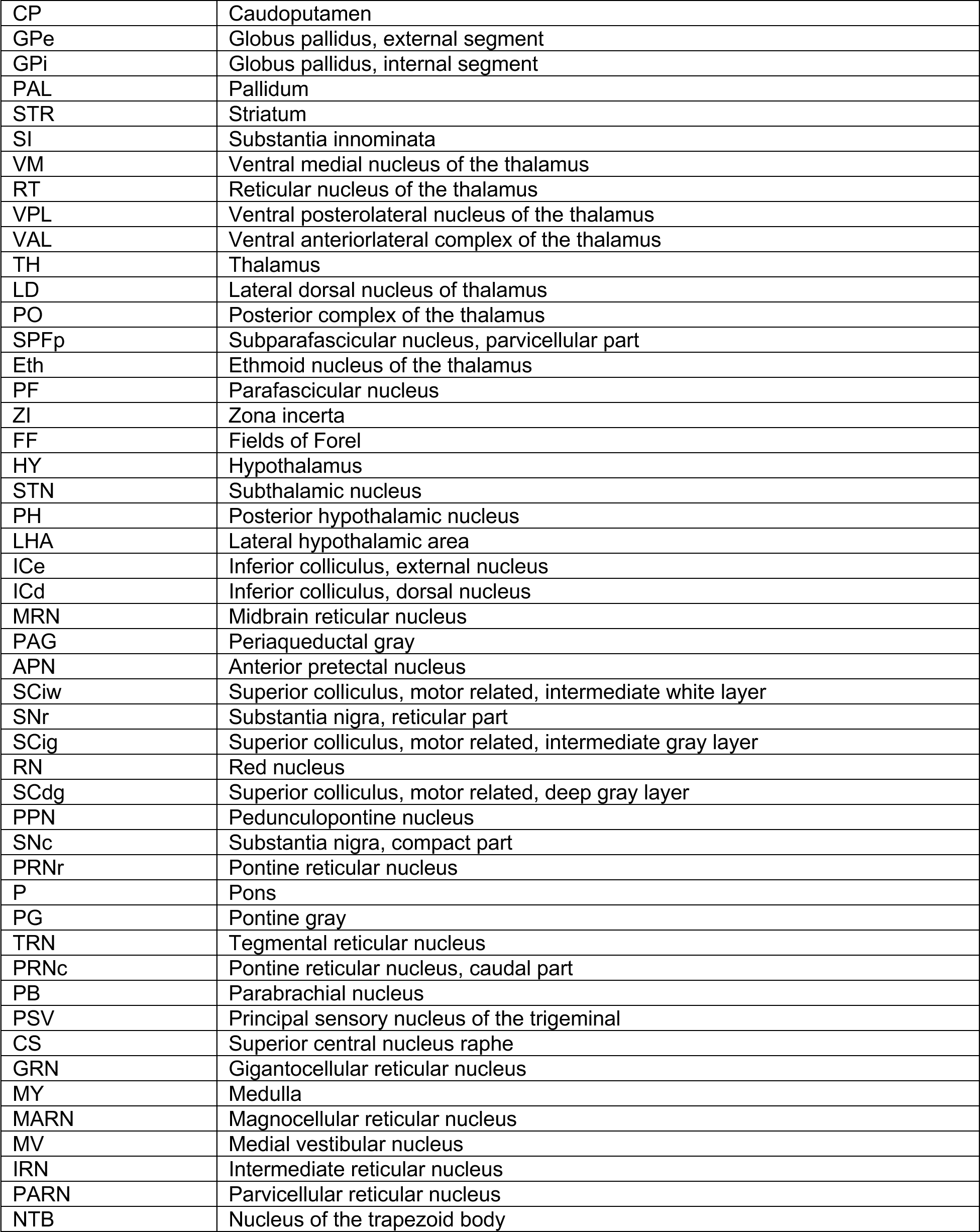

**Figure 3.**
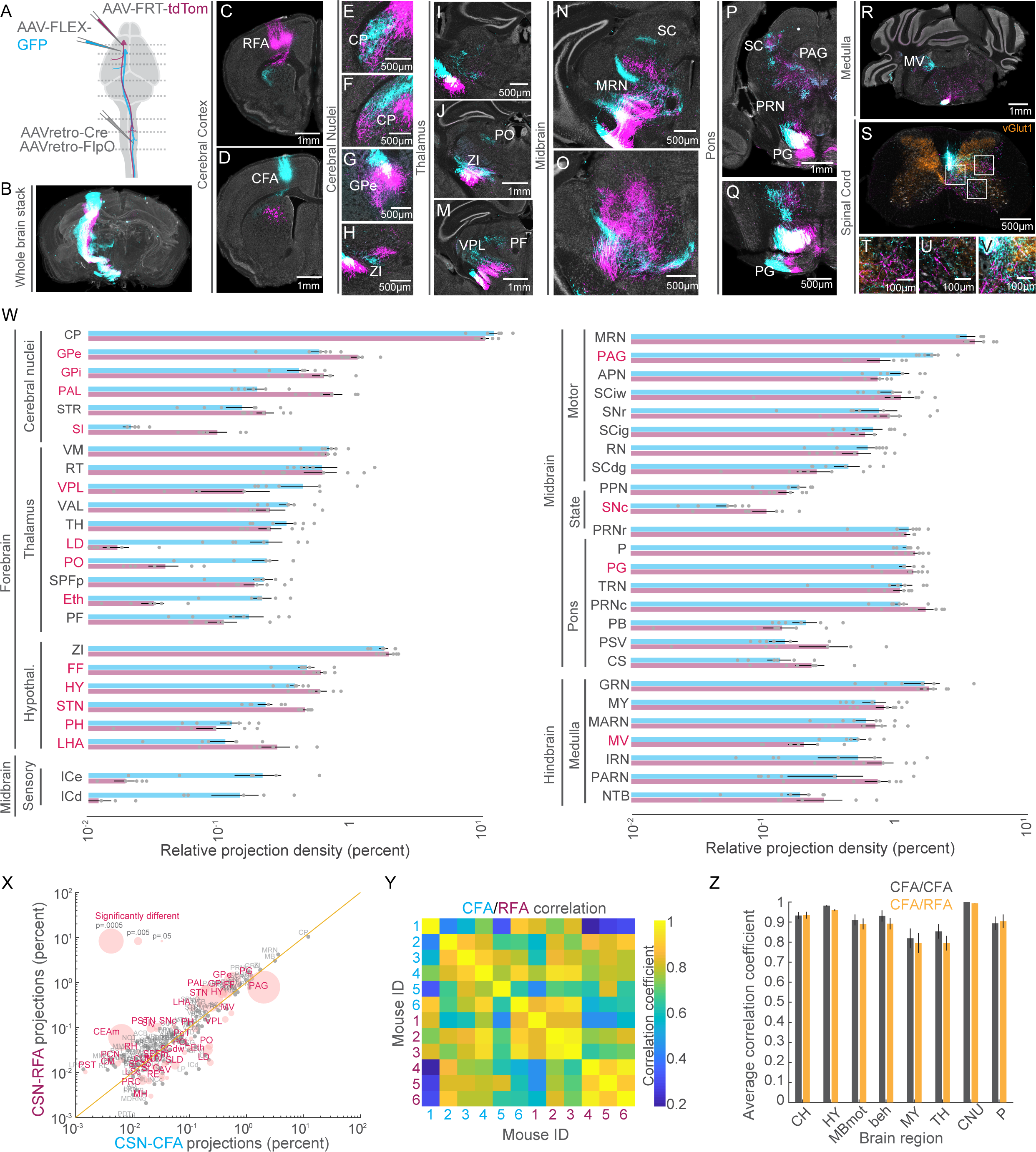
The supraspinal topography of CFA and RFA axon collaterals. **A)** Strategy to simultaneously label CFA and RFA CSNs. **B)** 3D reconstruction of a brain with CFA (cyan) and RFA (magenta) CSNs labeled. **C-V)** Exemplar photomicrographs of CFA and RFA CSN labeling throughout the central nervous system. Select regions are indicated. In S-V, vGlut1 axonal boutons are immunolabeled and shown in orange. **W)** The brain regions that are targeted by CFA and RFA CSNs, excluding isocortical structures. Regions are grouped by their ontology. Regions with significantly different fractions of CFA and RFA labeling are indicated in red. Note the log scale, given the large range of labeling. **X)** Scatter plot of regions containing CFA versus RFA projections. Regions with significantly different fractions of CFA and RFA labeling are colored red. The size of points corresponds to the p value of the comparison. Note the log scale, given the large range of labeling. **Y)** Matrix depicting the correlation of labeling within and across all 6 mice. Quadrants correspond to CFA versus CFA (upper left quadrant), RFA versus RFA (lower right quadrant), or CFA versus RFA (lower left and upper right quadrants) labeling. **Z)** The average correlation coefficients of CFA versus CFA (grey) labeling or CFA versus RFA labeling (orange) within major brain regions, across all mice. CH: cerebrum, HY: hypothalamus, MBmot: motor midbrain, beh: behavioral state related midbrain, MY: medulla, TH: thalamus, CNU: cerebral nuclei, P: pons. Refer to Table 3 for a complete list of the brain structures and their corresponding acronyms.

To determine the cell types that CSNs target at higher resolution, we mapped our complete dataset onto an atlas of spinal neuron diversity from Russ et al. comprising cross-validated RNA sequencing data compiled from several laboratories^5,30–33^. This harmonized atlas catalogues spinal cell type diversity at several levels of granularity, from broad neurotransmitter types to intersectional transcription factor expression that defines well-established spinal neuron subtypes (**Figure 2F, S1C,D**). We labeled our single nucleus transcriptomes for two classes by neurotransmitter type, glutamate for excitatory, and GABA and glycine for inhibitory (**Figure 2F** and **S1B**). First, we note a substantial preference for CSNs to contact inhibitory neurons over excitatory neurons (**Figure 2G**; 0.58±0.02 versus 0.23±0.06, p < 0.0001, Tukey’s multiple comparisons test). This result was obtained despite the relatively equal proportions of these two neuronal classes in the mouse spinal cord^34^. Next, we identified cell types by mapping their similarity to Russ’s 69 spinal neuron clusters that include 38 excitatory clusters and 27 inhibitory clusters^5^. Mapping to broad dorsoventral position of these clusters did not show a preference in the location of the populations targeted (**Figure 2H,I**). However, we found several specific spinal cell types targeted by CSNs, including excitatory and inhibitory neurons distributed across dorsal and ventral laminae (**Figure 2J**). Notably, approximately 25% of all neurons belonged to Inhibitory-23 cluster (27±2 %), likely corresponding to the Foxp2 clade of cardinal V1 interneurons and suggesting an Ia-inhibitory interneuron identity ^26^. This cluster was by a large margin the most prominent target of CSNs. Other prominent clusters of neurons included Excitatory-31 (9.07±1.12%), Inhibitory-12 (6.56±1.32%), Inhibitory-14 (5.88±0.94%), and Inhibitory-17 (8.26±1.15%), including cell types with well-described transcriptional profiles and lineages that indicate their functional identity^35,36^. Grouping these clusters into major spinal families, we identify ventral inhibitory and excitatory neurons as the most populous families of neurons targeted by CSNs (**Figure S1E,F**). Given that ventral neurons are classified into a single excitatory and inhibitory cluster each in this region because of their gradient of gene expression profiles, this was not an unexpected finding when considering the broad dorsoventral targeting of CSNs (VI: 30.48±2.22%, VE: 11.81±1.30%). This is followed by mid inhibitory (MI) neurons (12.12±1.53%), cerebrospinal fluid-contacting neurons (8.50±2.41%), Cdh3 neurons (7.01±1.33%), Chat neurons (5.84±0.94%), Npy neurons (6.60±1.30%), and Megf11 neurons (5.16±0.85%, **Figure S1F**). Together, these results map the connectivity between motor cortex and spinal neuron diversity, revealing well-defined cell types that predominate the neuronal targets of CSNs.

### CFA and RFA CSNs target partially distinct cell types

CSNs are anatomically and functionally diverse, raising the question of how innervation of spinal cell types differs across populations. We next determined the spinal cell types targeted by separate populations of CFA and RFA CSNs (**Figure 2K**). We first compared these subpopulations by estimating their broad dorsoventral location using the harmonized spinal neuron atlas. RNA sequencing of CFA_SC_ and RFA_SC_ populations significantly predicted a ventral horn identity for the majority of RFA_SC_ neurons, corroborating our anatomical tracing results and indicating RNA profiles alone may be used to predict spatial position (**Figure 2L, S2A**, RFA: dorsal 42.73±1.38% versus ventral 57.27±1.39%, p = 0.0003, unpaired t-test). We next measured the fractions of CFA_SC_ and RFA_SC_ neurons that belonged to each of the major spinal neuron families (**Figure S2B**). While CFA and RFA CSNs targeted mostly similar spinal cell types, MI neurons comprise a significantly larger fraction of CFA_SC_ neurons compared to RFA_SC_ neurons (**Figure 2M, S2C**, CFA: 15.37±1.68% versus RFA: 8.88±1.06%, p = 0.029, Sidak’s multiple comparisons test). We fractionated this comparison further, revealing cluster Inhibitory-17, which belongs to family MI, makes up a significantly larger fraction of CFA_SC_ neurons compared to RFA_SC_ neurons (**Figure S2D**, CFA: 10.80±1.01% versus RFA: 5.73±0.91%, p = 0.002, Sidak’s multiple comparisons test). This cluster, defined as deep dorsal horn neurons that co-express GABA and glycine, is implicated in sensorimotor processing and the presynaptic control of proprioceptive sensory afferents^5,36,37^. In addition, cluster comparison revealed Inhibitory-23 and Inhibitory-12 as more prominent targets of RFA CSNs (**Figure S2D**, Inhibitory-23, CFA: 24.64±3.71% versus RFA: 29.79±2.18%, p = 0.002; Inhibitory-12, CFA: 4.44±1.90% versus RFA: 8.69±1.21%, p = 0.02, Sidak’s multiple comparisons test). Cluster Inhibitory-23 is associated with type Ia inhibitory neurons of the V1 clade of ventral interneurons and is implicated in reciprocal inhibition^5,26^. These results show that CFA and RFA target mostly similar cell types in the spinal cord, but also identify biases in the spinal targets of CFA and RFA CSNs, uncovering a cell type-specific connectivity matrix.

### The supraspinal topography of CFA and RFA axon collaterals

CSNs control behavior through both direct projections to the spinal cord as well as collateral innervation of the brain^16,38^. The structures targeted by CSNs are widely distributed throughout the sensorimotor neuraxis, and the innervation of cell types in these regions is precisely maintained^7^. We next asked whether CFA and RFA CSNs differ in their brain targets, and if there are topographical features that distinguish innervation of individual brain structures. We used an intersectional strategy to specifically label CFA and RFA CSNs in the same mice. We injected AAV-retro-Cre and AAV-retro-FlpO in the cervical spinal cord, followed by AAV-FLEX-GFP into CFA and AAV-FRT-tdTomato into RFA (**Figure 3A**, same mice as Figure 1). We imaged antibody enhanced GFP and tdTomato labeling and again used BrainJ to register brain sections (**Figure 3B**). Visualizing GFP^+^ and tdTomato^+^ labeling revealed many brain regions targeted by CFA and RFA axon collaterals, spanning the full rostro-caudal extent of the central nervous system, including in the spinal cord (**Figure 3C-V**). We next mapped the positions of CFA and RFA CSN processes to a common brain atlas and ranked these brain regions by the fraction of total processes found in each structure (**Figure 3W**). We separated forebrain, midbrain, and hindbrain, and ranked the brain regions belonging to subregions of these major ontologies (i.e., substantia innominata belongs to cerebral nuclei belongs to forebrain). Consistent with previous studies, the caudoputamen (CP, or striatum) receives the single largest fraction of axonal collateralization (**Figure 1W**, CFA: 12.20±1.33%, RFA: 10.38±0.73%). We further compared the relative innervation of each region by CFA or RFA CSNs, leveraging the power of within-animal comparisons. While most brain regions receive indistinguishable fractions of CFA and RFA CSN input, we identified several brain regions that did differ in their relative innervation proportions. These include globus pallidus externus (**Figure 3G**, CFA: 0.570±0.090%, RFA: 1.119±0.124%, p = 0.0026, paired t-test), ventral posterolateral nucleus of the thalamus (**Figure 3M**, CFA: 0.424±0.146%, RFA: 0.151±0.086%, p = 0.018, paired t-test), and periaqueductal grey (**Figure 3P**, CFA: 2.008±0.243%, RFA: 0.789±0.160%, p = 7.14×10^−4^, paired t-test). The magnitudes of these differences vary substantially and are not clearly a function of the total fraction of projections (**Figure 3X**). Further, we independently rank ordered brain regions targeted by CFA or RFA, revealing the most densely innervated regions receive similar fractions of the total output of these populations (**Figure S3A-B**). Sorting RFA CSN projections by the rank order of CFA CSN projections also revealed similar fractions of CFA and RFA CSN projections were found in the top targeted brain regions, with some notable exceptions (**Figure S3C**).

We next sought to quantitate the similarity of labeling between CFA and RFA CSN collaterals across major brain regions (i.e., medulla, thalamus), as well as within condition reliability across mice. To this end, we created a matrix depicting the correlation coefficients of labeling across all mice (**Figure 3Y**). Visual inspection of the correlation matrix revealed no clear structure that would suggest major differences in innervation between CFA and RFA CSNs. We next measured the correlation of CFA and RFA CSN labeling split by major brain ontologies and compared these values to a control of CFA versus CFA innervation (**Figure 3Z**). Doing so we found no substantial differences in the innervation of major supraspinal structures, indicating brainwide innervation of CFA and RFA CSNs is similar. Still, we note that the terminal fields formed by CFA and RFA CSNs often appeared topographically distinct within individual brain regions, including caudoputamen (**Figure 3E-F**) and pontine grey (**Figure 3Q**). These results detail the collateral output structure of CFA and RFA CSNs, revealing largely similar innervation of the brain, but also some notable exceptions.

### CFA and RFA CSNs are defined by unique brainwide inputs

The information encoded in neuronal activity – and by extension neuronal function – is defined in part by synaptic inputs. Thus, revealing the inputs to CFA and RFA CSNs is essential to understanding their function. We harnessed the power of intersectional viral tracing to label inputs to CFA and RFA CSNs within the same animals. We first injected AAV-retro-Cre and AAV-retro-FlpO into cervical spinal cord. We then injected a 1:1 mixture of AAV-FLEX-N2cG and AAV-FLEX-TVA into RFA, followed by AAV-FLEX-N2cG and AAV-FRT-TVB in CFA. This led to the expression of rabies glycoprotein in CSNs and the expression of TVB and TVA in CFA and RFA CSNs, respectively. Two weeks later, we injected pseudotyped, G-deficient rabies viruses encoding fluorescent proteins into CFA and RFA. Specifically, we injected EnVA-N2cΔG-GFP in RFA and EnVB-N2cΔG-tdTomato in CFA (**Figure 4A**)^39^. This led to the expression of different fluorophores in inputs to CFA CSNs and RFA CSNs, all in the same animals. We used BrainJ to align sections and map neurons onto a common atlas (**Figure 4B**). Notably, the lack of reciprocal connectivity between CFA and RFA CSNs was reflected in the sparseness of dual-labeled CSNs (**Figure 4C-D**). This, along with a reported absence of long-range connectivity from CFA CSNs to RFA CSNs, supports the fact that GFP or RFP labeled inputs are not due to polysynaptic rabies transfer (i.e., jump across 2 synapses)^40^. Photomicrographs of neuronal labeling revealed widespread inputs to CFA and RFA CSNs, spanning much of the brain (**Figure 4E-T**). Simply with visual inspection, we observed major differences in the inputs to CFA and RFA CSNs. We quantified these differences by first independently ordering CFA or RFA inputs, revealing their distributions (**Figure S4A-B**). Sorting RFA labeling by the rank of inputs to CFA CSNs revealed a striking dissimilarity, indicating these two CSN populations receive substantially different input (**Figure S4C**).

**Figure 4.**
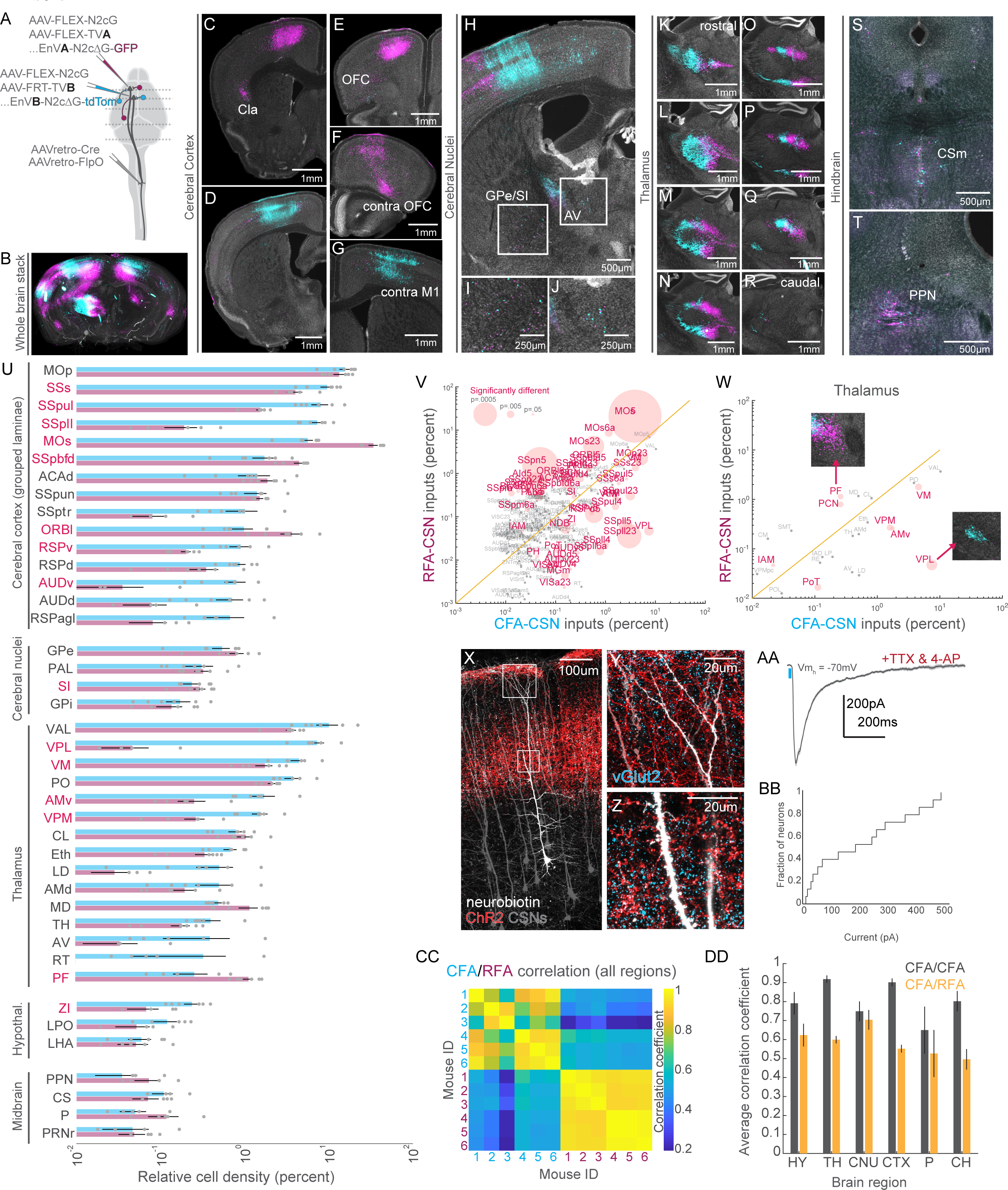
CFA and RFA CSNs are defined by unique brainwide inputs. **A)** Strategy to simultaneously label brainwide inputs to CFA and RFA CSNs. **B)** 3D reconstruction of a brain with inputs to CFA (cyan) and RFA (magenta) CSNs labeled**. C-T)** Exemplar photomicrographs of inputs to CFA and RFA CSN throughout the brain. Select regions are indicated**. U)** The brain regions that give rise to neurons that synapse on CFA or RFA CSNs. Regions are grouped by their ontology. Regions with significantly different fractions of input to CFA and RFA CSNs are indicated in red. Note the log scale, given the large range of labeling. **V)** Scatter plot of regions containing inputs to CFA versus RFA CSNs. Regions with significantly different fractions of CFA and RFA inputs are colored red. The size of points corresponds to the p value of the comparison. Note the log scale, given the large range of labeling. **W)** Same as **V),** but only showing thalamic inputs. The insets are photomicrophraphs illustrating the biases in thalamic input to CFA or RFA CSNs. **X)** Photomicrograph illustrating CSNs (grey), thalamocortical axons (red), and a select CSN targeted for whole cell, intracellular recording (white). **Y)** An expanded view of the superficial boxed region from **X)**, depicting axons surrounding the apical dendrites of CSNs. Presynaptic terminals are labeled with vGlut2 (cyan). **Z)** An expanded view of the lower boxed region from **X)**, depicting axons surrounding the trunk dendrites of CSNs. Presynaptic terminals are labeled with vGlut2 (cyan). **AA)** Example average EPSC evoked through optogenetic stimulation of thalamocortical axons (blue bar). Recordings are made in TTX and 4AP to isolate monosynaptic transmission. **BB)** Cumulative distribution of the current evoked through stimulation of thalamocortical axons. **CC)** Matrix depicting the correlation of input labeling within and across all 6 mice. Quadrants correspond to CFA versus CFA (upper left quadrant), RFA versus RFA (lower right quadrant), or CFA versus RFA (lower left and upper right quadrants) labeling. **DD)** The average correlation coefficients of inputs to CFA versus CFA (grey) or CFA versus RFA labeling (orange) within major brain regions, across all mice. Refer to Table 4 for a complete list of the brain structures and their corresponding acronyms.

**Table 4.**
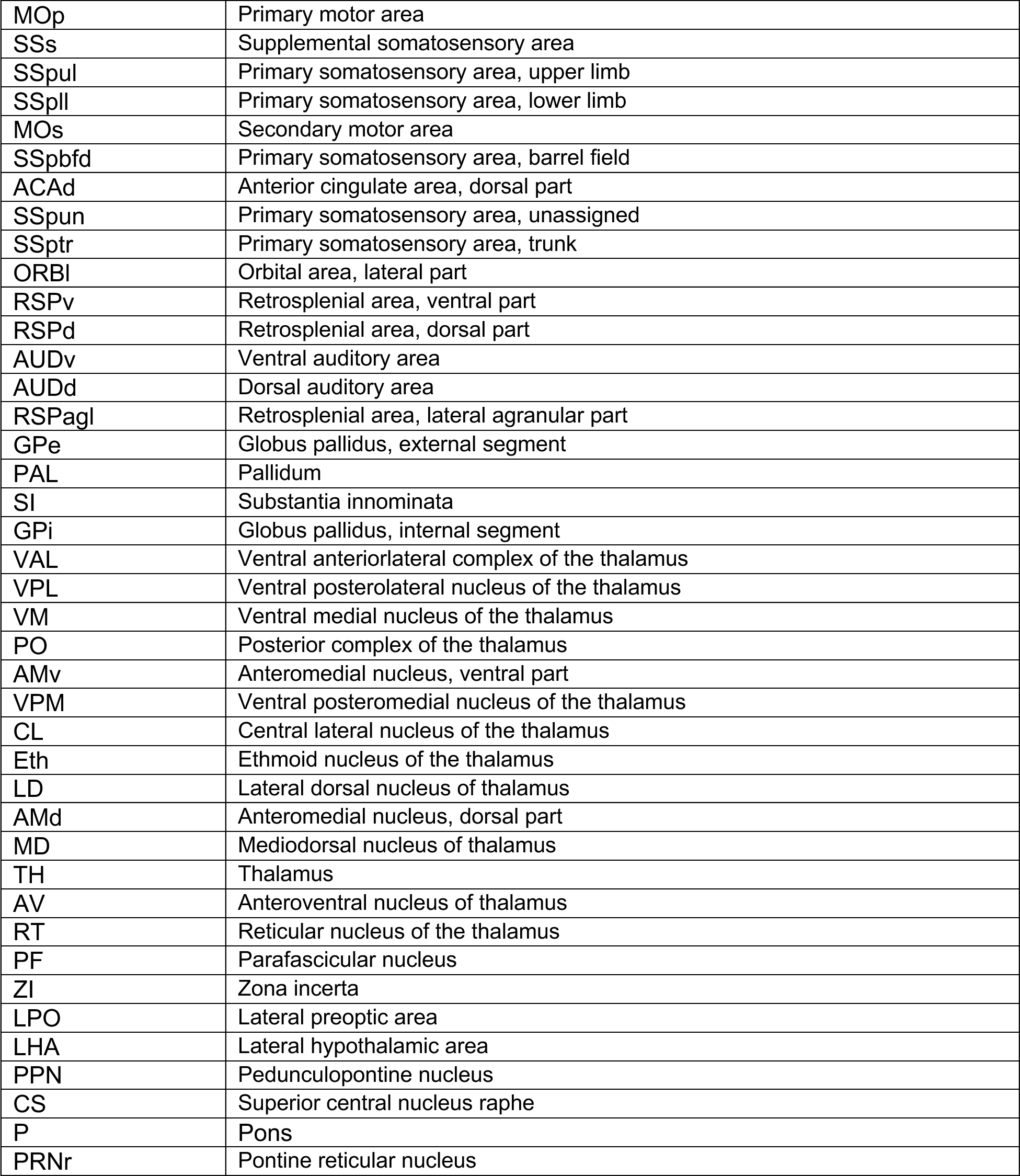

What are the different brain regions that provide synaptic input to CFA versus RFA CSNs? Quantification revealed that much of the input to all CSNs arises from isocortical structures, followed by substantial input from the thalamus (**Figure 4U-V**). Intriguingly, thalamic inputs to CFA or RFA CSNs are tessellated, a pattern easily appreciated in photomicrographs (**Figure 4K-R**). We observed a similar patchwork structure after CFA and RFA injections of dye-conjugated cholera toxins, which are transported both in retrograde and anterograde (**Figure S5A-B).** This pattern varies along the rostro-caudal axis of the thalamus, and maps onto well-defined thalamic regions, which we separately analyzed (**Figure 4W**). Of interest, CFA CSNs are strongly biased to receive inputs from ventral posterolateral thalamus among other regions, while RFA CSNs receive strong input from parafascicular thalamus (VPL: CFA: 7.43±1.55%, RFA: 0.047±0.027%, p = 0.005; PF: CFA: 0.258±0.114%, RFA: 1.139±0.159%, p = 0.019; paired t-tests).

Because thalamus has in the past been considered a non-canonical input to CSNs, we confirmed this connectivity using optogenetics-assisted electrophysiology. We expressed a fluorescent protein (FP) in CSNs by injecting AAV-retro-FP in the cervical spinal cord. We followed this with an injection of AAV-ChR2-FP into the caudal thalamus, resulting in the expression of ChR2 in thalamocortical axons. (**Figure 4X-Z, Figure S6A-B**). We then cut acute, live brain slices through motor cortex and made whole cell, patch clamp recordings from CSNs. While recording at membrane holding potentials to isolate excitatory currents, we optogenetically stimulated thalamic axons and recorded postsynaptic excitatory currents (EPSCs). ChR2 stimulation evoked EPSCs CSNs, even in the presence of TTX and 4AP to isolate monosynaptic transmission (**Figure 4AA-BB, Figure S6C-D**, 205.69±44.13pA, n = 15 cells, N = 3 mice). This confirmed thalamocorticospinal connectivity, and further reveals a driving synaptic force that mediates this connectivity.

Finally, we quantified the differences in brainwide input to CFA and RFA CSNs by measuring the correlation of labeling across mice and across brain regions, like our approach in Figure 3Y-Z. In stark contrast to axonal output labeling, inputs to CFA versus RFA CSNs were markedly uncorrelated (**Figure 4CC**). This result held true for several regions when we measured average correlation coefficients separated by major brain regions (**Figure 4DD**, HY: 0.791±0.038 v. 0.624±0.062, p = 0.0315; TH: 0.918±0.013 v. 0.599±0.017, p = 0.0000; CNU: 0.749±0.033 v. 0.704±0.041, p = 0.2926; CTX: 0.901±0.013 v. 0.552±0.020, p = 0.0000; P: 0.650±0.078 v. 0.526±0.066, p = 0.1910; CH: 0.802±0.034 v. 0.496±0.030, p = 0.0000). Nearly all major brain ontologies with substantial input to CSNs differ in the substructures that target CFA versus RFA CSNs. These results reveal CFA and RFA CSNs receive substantially different brainwide inputs, more so than their outputs, and this connectivity principle is what likely determines differential activity profiles and behavioral functions of these populations.

## DISCUSSION

This study reveals both the regional and cellular spinal targets of two major populations of CSNs, as well as the input and output organization of these populations in the brain. It is first worth considering the known differences in structure and function between CFA and RFA. Their rostro-caudal separation has led to the suggestion that RFA may be involved in motor planning, and CFA more involved in execution. However, microstimulation, recording, and manipulation studies found mixed results, suggesting preparation and execution are encoded in both regions, while others concluded CFA and RFA are distinct somatotopic representations of the forelimb^8,14,19,41^. One obstacle for synthesizing these results is the difference in methods used and specific circuits studied. For one, motor cortex is incredibly diverse in cellular organization, confounding the interpretation of recordings and manipulations of large populations of cortical neurons^42^. Even activation and inactivation of isolated cell types, including CSNs, influences activity in brain regions innervated by their collaterals^43^. Our approach here has been to first chart the logic of CFA and RFA CSN connectivity, so that later studies can perform recordings and manipulations informed by the circuit architecture.

We first note that CSNs formed synapses on dispersed groups of spinal neurons that span many functionally distinct laminae. Despite this widespread innervation, AnteroT-seq revealed CSNs are remarkably selective in the spinal neuron types they targeted, with a quarter of all CSN_SC_ neurons belonging to a single spinal neuron cluster. Inhibitory neurons made up a large majority of CSN targets, which could reflect a prominent role for CSNs in suppressing movement or sensory feedback. Alternatively, this bias may prevent overexcitability through feedforward inhibition. In addition, CFA and RFA CSNs largely targeted similar cell types, despite the spatial separation of these populations in sensorimotor cortex. Still, there are notable exceptions to this general trend. For example, we found a significantly larger proportion of CFA_SC_ neurons compared to RFA_SC_ neurons in the deep dorsal horn family MI and its cluster Inhibitory-17, neurons thought to be involved in suppressing proprioceptive afferents^36,44^. This suggests that CFA CSNs, or a fraction of that population, is more involved in the regulation of sensory feedback during movement compared to RFA CSNs, perhaps constituting a corollary discharge circuit^45^. Conversely, our finding that RFA CSNs targeted deeper spinal neurons indicates these neurons may have privileged access to spinal circuits closer in location – if not function – to motor neurons. Along those lines, more RFA_SC_ neurons than CFA_SC_ neurons belonged to the ventral inhibitory cluster Inhibitory-23, a glycinergic population of putative V1 lineage. Perhaps RFA CSNs play a larger role in gating reciprocal inhibition of motor pools through synapses on Ia interneurons born of V1 lineage^46^. The select recruitment of these corticospinal populations could allow for flexible alternation and co-contraction of antagonist muscles, a behavioral switch that requires co-variation of motor cortex activity^47^.

In alignment with our previous study, we identify the striatum as receiving the single largest fraction of supraspinal corticospinal axon collaterals^7^. Overall, CFA and RFA CSNs formed axon collaterals in the brain that were similar in their major targets, so that the information encoded in these populations may be uniformly broadcast through the brain. Still, CFA and RFA CSN may target different cell types or subregions of these brain structures. Future brainwide transsynaptic sequencing and optogenetics-assisted circuit mapping efforts will shed light on this possibility. In stark contrast to their outputs, we identified highly divergent brainwide inputs to CFA and RFA CSNs, a result that implies the information encoded across populations differs. Much of the divergent input arose from isocortex and thalamus, rather than midbrain and hindbrain structures. This raises the possibility that CSNs are integrating different higher order associative information, rather than different low-level state information that would be encoded in ascending brainstem regions, where inputs were intermingled.

Outside of CFA and RFA, there are several other motor and sensory cortical regions that contain CSNs, including neurons that innervate hindlimb lumbar regions of spinal cord^48–50^. These hindlimb cortical regions are important for behaviors including obstacle avoidance during locomotion^48^. It will be interesting to extend our level of cellular and circuit characterization to all these distinct populations to compare the spinal neurons differentially targeted. Finally, CSN populations act to broadcast neuronal activity throughout the brain and spinal cord. This activity is complex, task-dependent and affects behavior through the neural circuits that CSNs target^7,51–53^. Our study identifies the structures and cell types distributed throughout the nervous system that receive input from two essential corticospinal populations, revealing unique circuitry that dictates the influence of descending signals on motor output and sensory feedback.

## ACKNOWLEDGEMENTS

We are grateful to L. Hammond and F. Fiederling for creating exceptional brain and spinal cord image analysis pipelines, as well as helping with these tools. We thank I. Shieren and the Zuckerman Institute’s Flow Cytometry Platform for assistance with FACS. We thank S. Brenner-Morton for antibodies. Imaging was performed in part with support from the Zuckerman Cellular Imaging platform at Columbia University and the National Institute of Health (NIH 1S10OD023587-01). CVS-N2c rabies viruses were produced by the Center for Neuroanatomy with Neurotropic Viruses, supported by P40 OD010996. A.N. is supported by an NIH Pathway to Independence Award (R00NS118053). L.M.C is supported by a NIH Pathway to Independence Award (K99NS127857). R.M.C. is supported by 5U19NS104649 and the Allen Institute for Brain Science.

## AUTHOR CONTRIBUTIONS

L.M.C, R.M.C. and A.N. designed experiments. L.M.C. and A.N. performed, analyzed, and interpreted experiments. E.T.T, K.S., and B.T. performed RNA sequencing. A.N. wrote the original draft of the manuscript with edits from L.M.C., B.T. and R.M.C.

## MATERIALS AND METHODS

### Experimental model and animals

Adult mice of both sexes, aged 2-6 months, were used for all experiments. The strains used were C57BL6/J (Jackson Laboratories, 000664) and B6;129-Gt(ROSA)26Sort^m5(CAG-Sun1/sfGFP)Nat/^J (CAG-Sun1/sfGFP, Jackson Laboratories, 021039). All mice were kept under a 12-h light-dark cycle.

### Stereotaxic viral injections

Analgesia in the form of subcutaneous injection of carprofen (5 mg per kg body weight) was administered on the day of the surgery, along with bupivacaine (2 mg per kg body weight). Mice were anesthetized with isoflurane and placed in a stereotaxic holder (Kopf). A midline incision was made to expose the skull, and a craniotomy was made over the injection site. A pulled glass pipette was filled with virus, and a Nanoject III was used to make multiple small-volume injections across into the spinal cord, with parameters that depended on the experiment and reagents used. To label CFA and RFA CSNs, 50nL each of AAV-FLEX-GFP and AAV-FRT-tdTomato were injected into CFA and RFA, respectively. Each injection was targeted to one location, approximately 700um below the pia. To label spinal neurons targeted by CFA and RFA, 400-600nL of AAV1-Cre and AAV1-FlpO were injected into CFA and RFA, respectively. To label spinal nuclei for sequencing, 400-600nL of AAV1-Cre and AAV1-FlpO were injected bilaterally into CFA and RFA, respectively. For optogenetics-assisted electrophysiology, 50nL of AAV1-ChR2-GFP was injected into the thalamus, centered between PO and PF. To label inputs to CFA and RFA CSNs, the following strategy was used. Into mice that express Cre and FlpO in CSNs (see below), 50nL total of a 1:1 mixture of AAV-FLEX-N2cG and AAV-FLEX-TVA.mCherry was injected into RFA. 50nL of a 1:1 mixture of AAV-FLEX-N2cG and AAV-FRT-TVB was injected into CFA. After two weeks, 300nL of EnVA-N2cΔG-GFP was injected into RFA and 300nL of EnVB-N2cΔG-tdTomato was injected into CFA.

### Spinal cord viral injections

Analgesia in the form of subcutaneous injection of carprofen (5 mg per kg body weight) was administered the day of the surgery, along with bupivacaine (2 mg per kg body weight). Mice were anesthetized with isoflurane and placed in a stereotaxic holder (Kopf). A midline incision was made to expose the spinal column. The musculature overlying the column was resected, and the T2 process was secured to minimize spinal cord movement. The tail was gently stretched with another spinal clamp to separate the vertebrae. A surgical microknife and fine forceps were used to sever the meninges, exposing the spinal cord. A pulled glass pipette was filled with virus, and a Nanoject III was used to make multiple small-volume injections across into the spinal cord, with parameters that depended on the experiment and reagents used. To label inputs to the spinal cord, 200nL of AAV-retro-mCherry was injected into each segment C3-C8 of the cervical spinal cord. Injections were split across two separate penetrations spanning the mediolateral extent of the spinal grey. Within each penetration the injection was spread across the entire superficial to ventral extent of the spinal cord. To label CFA and RFA CSNs, 200nL of a 1:1 mixture of AAV-retro-Cre and AAV-retro-FlpO was similarly injected into C3-C8 of the cervical spinal cord. To label spinal neurons targeted by CFA and RFA in the same mice, 200nL of a 1:1 mixture of AAV-FLEX-H2b-GFP and AAV-FRT-H2b-RFP was injected into C3-C8 of the cervical spinal cord. To label inputs to CFA and RFA CSNs, 200nL of a 1:1 mixture of AAV-retro-Cre and AAV-retro-FlpO was injected into C3-C8 of the cervical spinal cord. Following all injections, the skin was sutured closed, and animals were closely monitored during recovery.

### Nuclear suspension and FACS

Six weeks post injection, mice were deeply anesthetized with isoflurane and transcardially perfused with a carbogenated, ice-cold artificial cerebrospinal fluid (ACSF) solution containing: 100mM NaCl, 10mM HEPES, 25mM Glucose, 75mM Sucrose, 7.5mM MgCl_2_, and 2.5mM KCl. A ventral laminectomy was performed, and the cervical spinal cord was rapidly isolated in carbogenated, ice-cold ACSF. The meninges were carefully removed with Vannas scissors and forceps, and the cervical spinal cord was divided into hemisections and rapidly frozen on dry ice. Samples were stored at −80C until the day of nuclear isolation. Nuclei were isolated using the 10x Chromium Nuclei Isolation Kit following the manufacturer protocol. All solutions were made fresh from each prep. DAPI was added to the samples after the final resuspension, and samples were filtered through a 30 um filter (CellTrics, Sysmex). Fluorescence Activated Cell Sorting was performed immediately after dissociation at the Zuckerman Institute Flow Cytometry platform using a MoFlo Astrios EQ (Beckman Coulter) sorter. Sorting was performed with 100 um nozzle at sheath pressure of 28 PSI. Sample pressure was maintained below 28.5 PSI. The following laser lines and filters were used: For DAPI the 205 nm laser was used with a 448/59 band-pass filter. The PMT was set at 315V with an amplifier gain of 2. For GFP, the 488 nm laser was used with a 513/26 band-pass filter. The PMT was set at 370V with an amplifier gain of 2. These fluorochromes required different laser sources on different laser paths which enabled very clean separation of the emission fluorescence and no need for fluorescence compensation. The 488 laser was also used for detection of forward and side scatter. Samples were first gated of forward and side scatter to eliminate debris, electronic noise, and large particles. Doublet discrimination was performed using a combination of data plots consisting of Side Scatter height vs. Side Scatter Area, Side Scatter height vs. Side Scatter width, and DAPI Log height vs. DAPI width. DAPI and GFP double positive nuclei were sorted using “Single Cell” mode with a drop envelope of 1. Nuclei were collected in 8 well PCR strips preloaded with lysis solution. Both the cell suspension and the collection chamber were maintained cold during sorting. PCR strips with collected nuclei were spun down immediately after sorting and frozen on dry ice.

### Single-nucleus RNA-sequencing

#### RNA amplification, library preparation, and RNA sequencing

SMART-Seq v4 Ultra Low Input RNA Kit for Sequencing (Takara #634440) was used per manufacturer’s instructions for cDNA synthesis of single cell RNA and subsequent amplification, with the exception that all reaction volumes were reduced to 0.5x. Single cells were stored in 8-strips at −80°C in 5.25 μl of collection buffer (SMART-Seq v4 lysis buffer at 0.83x, RNase Inhibitor at 0.17 U/μl, and ERCC MIX1 at final 1×10-8 dilution as described above). Twelve to 36 8-strips were processed at a time (the equivalent of 1-3 96-well plates). At least 1 control strip was used per amplification set, containing 2 wells without cells (termed ERCC), 2 wells without cells or ERCC (termed NTC), and 2 wells of 10 pg of Mouse Whole Brain Total RNA (Zyagen, MR-201) and 2 wells of 10 pg Control RNA provided in the Takara kit.

Mouse nuclei were subjected to 22 or 23 PCR cycles after the reverse transcription step. SPRI bead (Sera-Mag Select beads GE Healthcare #29343057) purification was done using the Agilent Bravo NGS Option A instrument. A bead ratio of 1x was used (25 μl of Sera-Mag Select beads to 25 μl cDNA PCR product with 0.5 μl of 10x lysis buffer added, as per Takara instructions at 0.5x volume), and purified cDNA was eluted in 17 μl elution buffer provided by Takara. All samples were quantitated using PicoGreen® on Molecular Dynamics M2 SpectraMax instrument. The samples were then run on the Agilent Fragment Analyzer (96) using the High Sensitivity NGS Fragment Analysis Kit (1bp-6000bp) to qualify cDNA size distribution. Purified cDNA was stored in 96-well plates at −20°C until library preparation.

All samples proceeded through NexteraXT DNA Library Preparation (Illumina FC131-1096) using custom 8-base Unique Design Index primers designed and manufactured by IDT (Integrated DNA Technologies). NexteraXT DNA Library prep was done at 0.2x volume on the Mantis instrument (Formulatrix). Reduction in volume was applied to input and all reagents, but otherwise the manufacturer’s instructions were followed. 50pg cDNA input was used in generating the libraries. An aliquot of all amplified cDNA samples was first normalized to 50 pg/ul with Nuclease-Free Water (Ambion), then this normalized sample aliquot was used as input material into the NexteraXT DNA Library Prep (for a total of 50pg). Before bead purification after PCR, 12.1ul of Nuclease-Free Water (Ambion) was added to the 10ul PCR reaction volume, bringing the total volume to 22.1ul post-PCR and before bead purification. SPRI Sera-Mag Select bead purification was done using the Agilent Bravo NGS Option A instrument. A bead ratio of 0.9x was used (20 ul of Sera-Mag Select beads to 22.10 ul library product, as per Illumina protocol), and all samples were eluted in 22 μl of Resuspension Buffer (Illumina). All samples were quantitated using PicoGreen using Molecular Bynamics M2 SpectraMax instrument. All samples were run on the Agilent Fragment Analyzer (96) using the High Sensitivity NGS Fragment Analysis Kit (1bp-6000bp) for sizing. Molarity was calculated for each sample using average size as reported by the Fragment Analyzer and pg/μl concentration as determined by PicoGreen. Samples (5 μl aliquot) were normalized to 2-5 nM with Nuclease-free Water (Ambion), then 2 μl from each sample within one 96-index set was pooled to a total of 192 μl at 2-5 nM concentration. Libraries were further multiplexed at 768 samples/flowcell by pooling 8 96-sample libraries using compatible Index Sets. A portion of the final library pool was sequenced on an Illumina NextSeq2000 instrument using the P2 flowcell, for a target of 500,000 reads per single cell or nucleus.

#### Data processing of RNA sequencing

RNA-Seq alignment and data processing was done the same way for all single cell and single nucleus RNA-Seq data, except that different versions of the genome and transcriptome were used for each species.

The raw fastq files from Illumina were trimmed using the fastqMCF program (Aronesty, et al., 2011) The trimmed paired-end reads were mapped and counts generated using STAR aligner (v2.7.1a) with default settings against mouse mm10 GENCODE vM23/Ensembl 98 reference genome, downloaded from 10X cell ranger (refdata-cellranger-arc-mm10-2020-A-2.0.0). STAR uses and builds its own suffix array index which considerably accelerates the alignment step while improving on sensitivity and specificity, due to its identification of alternative splice junctions. Next, the duplicates were removed using STAR as well. Reads that did not map to the genome were then aligned to synthetic constructs (i.e. ERCC) sequences and the E.coli genome (version ASM584v2). The final results files included quantification of the uniquely mapped reads (raw exon and intron counts for the transcriptome-mapped reads). Included in the final results files are the percentages of reads mapped to the transcriptome, to ERCC spike-in controls, and to E.coli.

#### QC and analysis of RNA sequencing

Nuclei with <1000 genes, <100,000 total read, <75% aligned reads, >0.5 CG complexity, and >5% mitochondrial RNA were excluded from further analysis. All analysis was performed using Seurat. Mapping to data from Russ, *et al* was also performed with Seurat using transfer anchors.

### Slice electrophysiology

Mice were deeply anesthetized with isoflurane and transcardially perfused with an ice-cold carbogenated high-magnesium (10 mM) artificial cerebrospinal fluid (ACSF). The brain was removed from the skull and glued to the stage of vibrating microtome (Leica). 300 µm coronal brain slices were cut in a bath of ice-cold, slushy, carbogenated low-calcium ACSF. Slices were immediately transferred to a 37C bath of normal ACSF containing 124 mM NaCl, 2.7 mM KCl, 2 mM CaCl2, 1.3 mM MgSO4, 26 mM NaHCO3, 1.25 mM NaH2PO4, 18 mM glucose and 0.79 mM sodium ascorbate, where they incubated for 25 minutes. Slices were then moved to room temperature, where they remained for the duration of the experiment. Patch electrodes (2–6 MΩ) were filled with a potassium gluconate-based internal solution containing 135 mM potassium gluconate, 2 mM MgCl2, 0.5 mM EGTA, 2 mM magnesium ATP, 0.5 mM sodium GTP, 10 mM HEPES, 10 mM phosphocreatine and 0.15% Neurobiotin. All recordings were made using a Multiclamp 700B amplifier, the output of which was digitized at 10 kHz (Digidata 1440A). Series resistance was always <30 MΩ and was compensated up to 90%. Neurons were targeted with DIC microscopy and epifluorescence when appropriate. In a subset of experiments, cell morphology was visualized through internal dialysis of 0.1 mM Alexa Fluor 594 cadaverine or 0.1 mM Alexa Fluor 488 sodium salt. ChR2-expressing axons were photostimulated using 10-ms pulses of 473-nm LED light (CoolLED) delivered through a 10x objective centered over the recording site. Brain slices were histologically processed to visualize Neurobiotin-filled cells through streptavidin-Alexa Fluor processing.

### Histology and confocal imaging

Mice were anesthetized with isoflurane and transcardially perfused with ice cold PBS followed by cold 4% paraformaldehyde. Brains and spinal cords were fixed overnight in 4% paraformaldehyde and cryopreserved in a 30% sucrose solution at 4°C for 3 days or until they sunk. Brains were embedded in Optimum Cutting Temperature Compound (OCT, Tissue-Tek) and frozen on dry ice. Spinal cords were prepared for SpinalJ processing by mounting them in dissolvable SpineRacks embedded within OCT. The spinal cord was cut into three segments comprising the cervical, thoracic, and lumbar regions. Each of these regions was further divided into 3 chunks, and these chunks were oriented rostral side downward in a 3×3 array SpineRack submerged in OCT within a mold. After SpineRacks had fully softened in OCT, they were frozen on dry ice. Sections between 50 µm and 80 µm were cut using a cryostat (Leica). Spinal cord sections were directly mounted on slides, while brain sections were deposited in 24 well plates. Tissue was rinsed several times in PBS, and permeabilized in 0.2% Triton X-100 (PBST). Immunostaining was performed with primary antibodies diluted to working concentration for 3 days at 4°C, and with secondary antibodies (Jackson ImmunoResearch) diluted 1:1000 overnight at 4°C. Brain and spinal cord slices mounted to slides were briefly incubated with TrueBlack diluted in 70% ethanol to quench lipofuscin and background autofluorescence. Confocal imaging was performed on a Zeiss 710 or Zeiss 880 using a 10x, 20x, or 40x objective.

### Slide scanning and anatomical reconstructions

Sections were imaged using an AZ100 automated slide scanning microscope equipped with a 4x 0.4-NA objective (Nikon). Image processing and analysis using BrainJ or SpinalJ was as previously described. Briefly, a seven-pixel rolling-ball filter was used on all images to reduce background signal and a machine-learning pixel classification approach using Ilastik was used to identify cell bodies and neuronal processes. To map the location of these structures to an annotated brain or spinal cord atlas, 3D image registration was performed using Elastix relative to a reference brain or spinal cord. The coordinates of detected cells and processes were then projected into the Allen Brain Atlas Common Coordinate Framework. Visualizations of the data were performed in ImageJ, and subsequent analyses were performed in MATLAB using custom software.

### Statistics and data analysis

Statistical tests and significance are reported throughout the manuscript. Statistical tests were performed in MATLAB and Prism. Significance is defined as p < 0.05.

---

**Figure S1.**
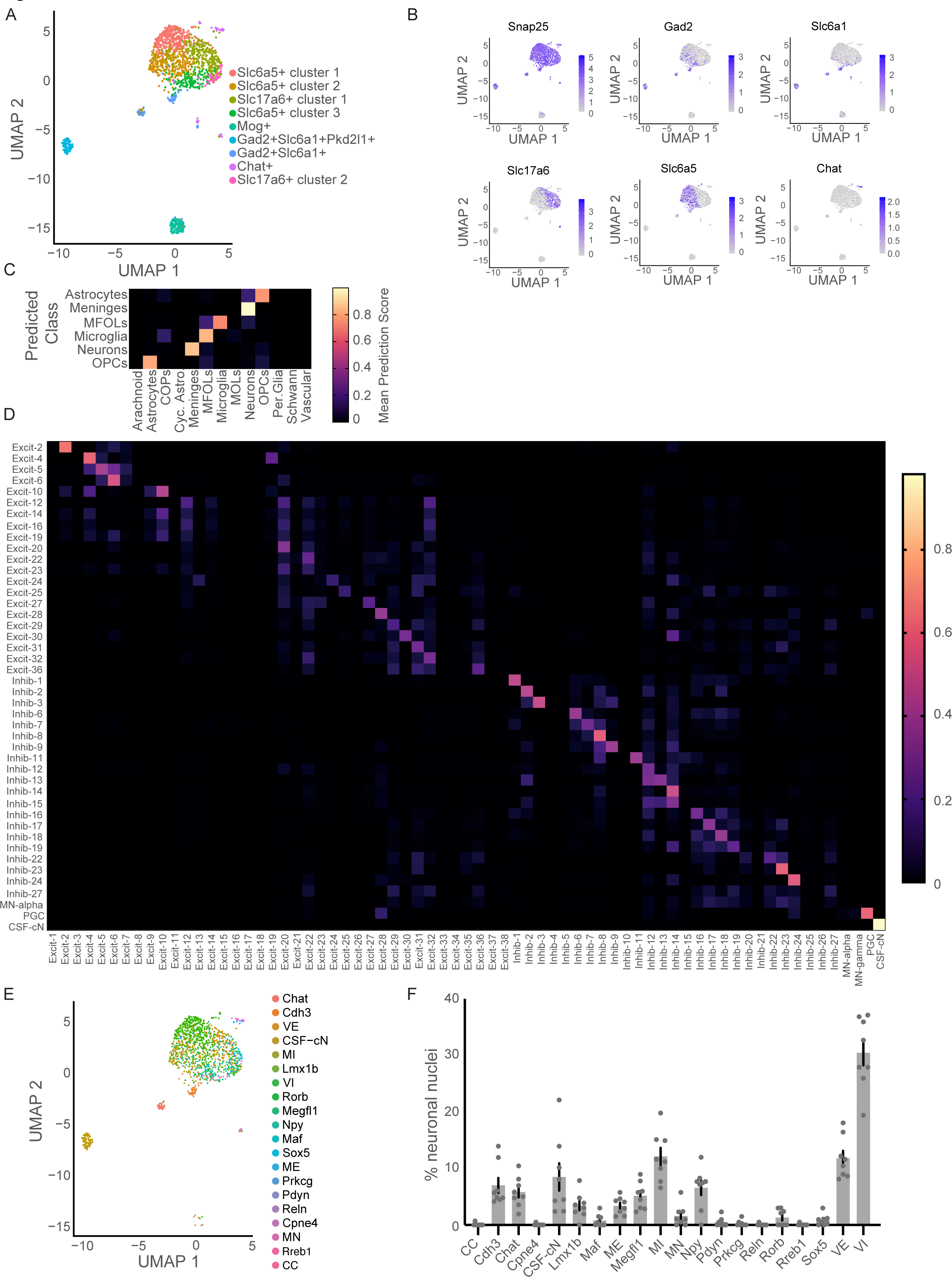
**A)** UMAP plot of all CSN_SC_ cells, labeled with factors that define cluster identity. **B)** UMAP plots depicting the expression of various factors. **C)** Heatmap of the average prediction scores for major neuron types captured. Scores for all types are included but not all types are present in data. **D)** Heatmap of the average prediction scores for the identified clusters. Scores for all clusters are included but not all clusters are present in data. **E)** UMAP plot of all CSN_SC_ neurons, labeled by major spinal neuron family. **F)** The percentage of neuronal nuclei belonging to each major family, across individual experiments. Error bars indicate SEM.

**Figure S2.**
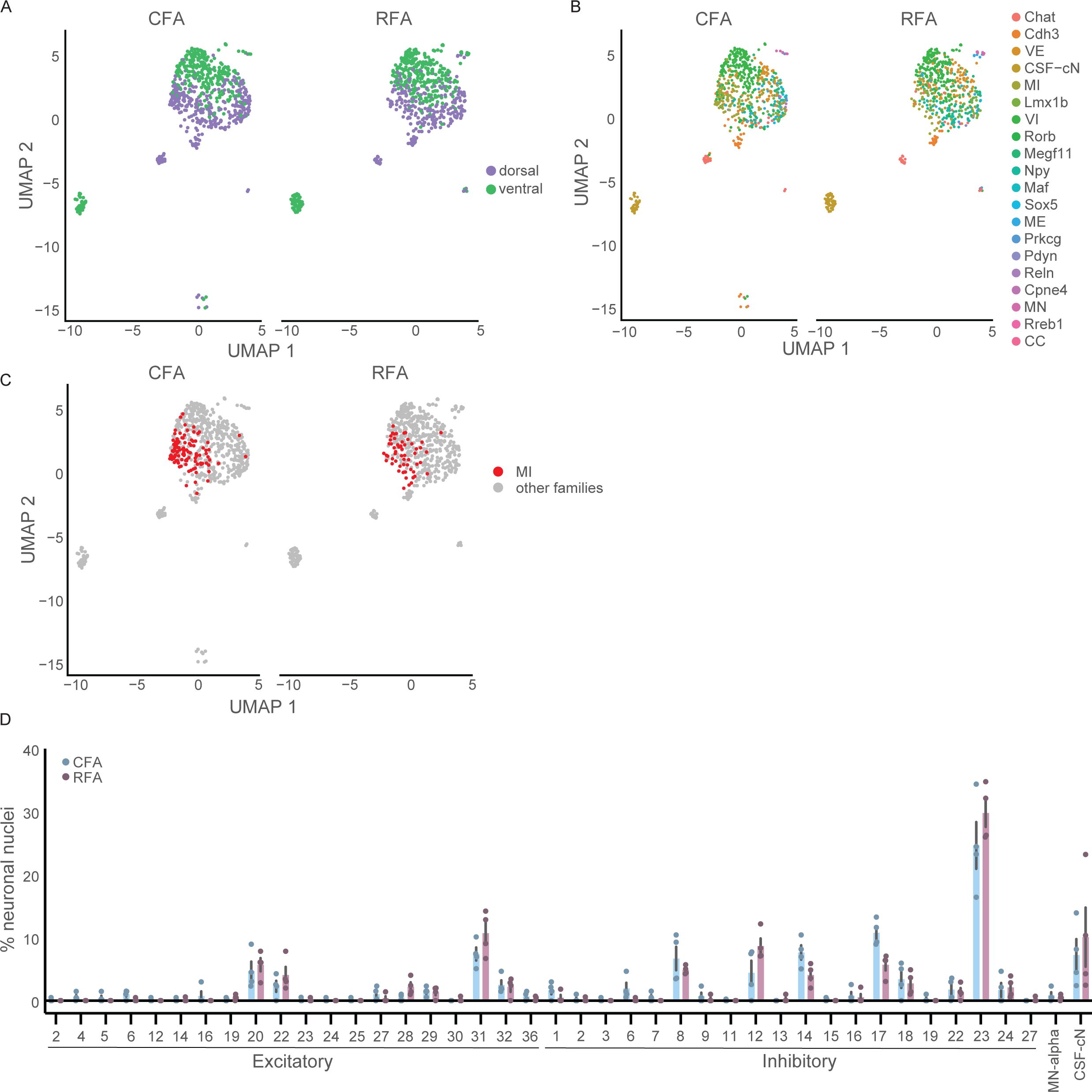
**A)** UMAP plots of CFA_SC_ neuronal nuclei, separated into CFA_SC_ and RFA_SC_ conditions. **B)** UMAP plots of CFA_SC_ and RFA_SC_ neuronal nuclei, labeled by major spinal neuron family. **C)** UMAP plots of CFA_SC_ and RFA_SC_ neuronal nuclei, with MI-labeled nuclei colored in red. **D)** The percentage of CFA_SC_ and RFA_SC_ neuronal nuclei belonging to each identified cluster, across individual experiments. Error bars indicate SEM.

**Figure S3.**
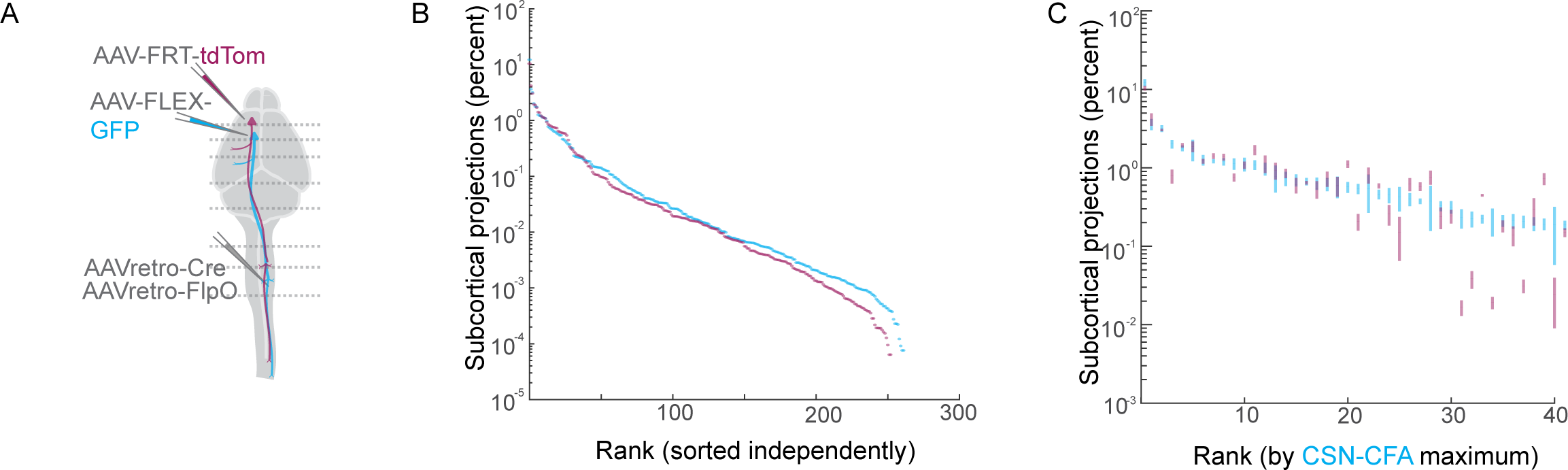
**A)** Review of the anatomical tracing strategy. **B)** All brain regions, independently ranked by the fraction of either CFA (blue) or RFA (magenta) CSN projections found in each structure. **C**) The top 40 brain regions targeted by CFA CSN projections (cyan), as well as RFA CSN projections sorted by the same order. The height of the vertical bar reveals the SEM of the projections in each structure.

**Figure S4.**
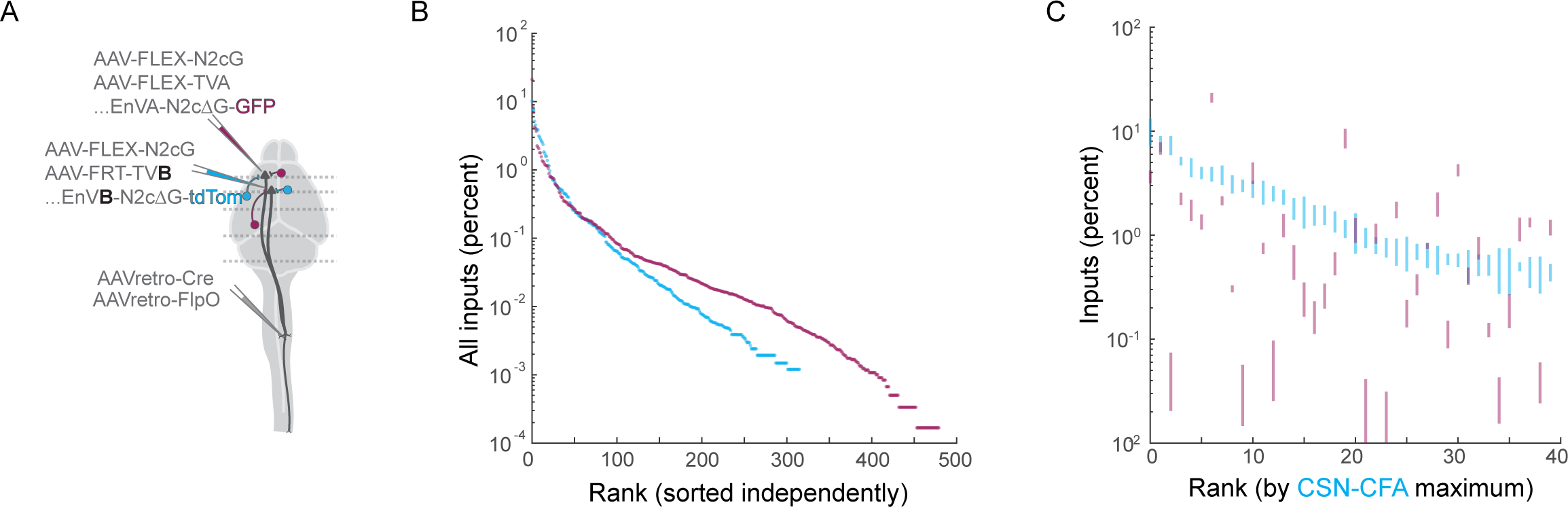
**A)** Review of the anatomical tracing strategy. **B)** All brain regions, independently ranked by the fraction of inputs to either CFA (blue) or RFA (magenta) CSNs found in each structure. **C**) The top 40 brain regions synapsing on CFA CSNs (cyan), as well as RFA CSNs sorted by the same order. The height of the vertical bar reveals the SEM of the projections in each structure.

**Figure S5.**
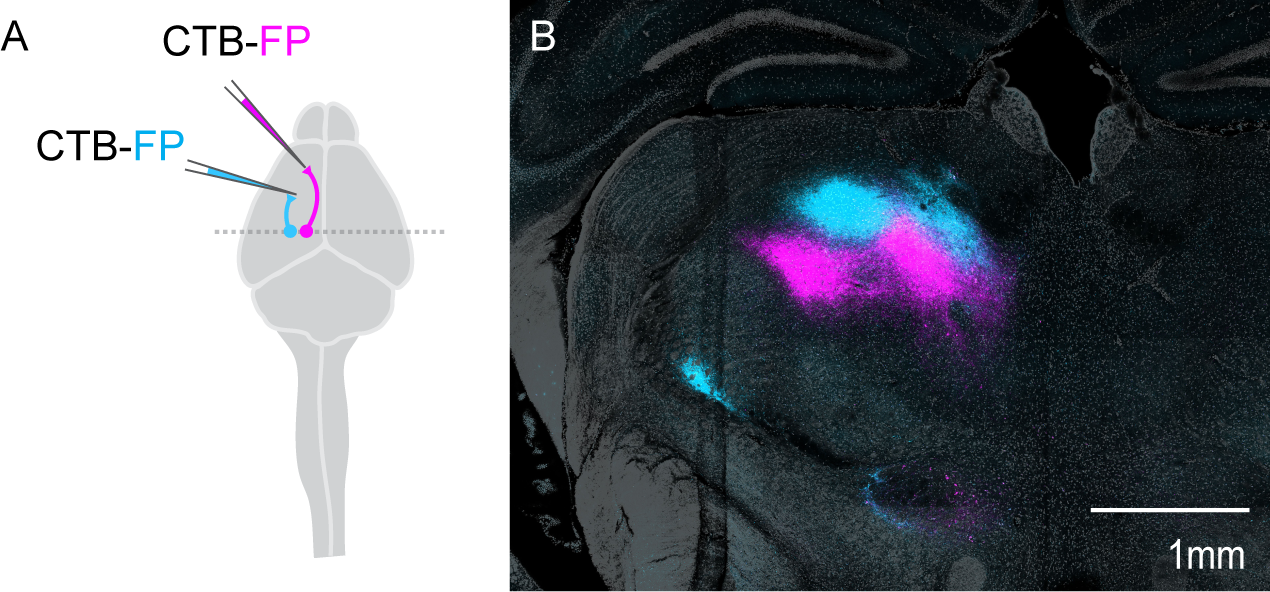
**A)** Strategy to label inputs to CFA and RFA using conventional tracers. **B)** Photomicrograph of the thalamus depicting structures connected to CFA (cyan) or RFA (magenta).

**Figure S6.**
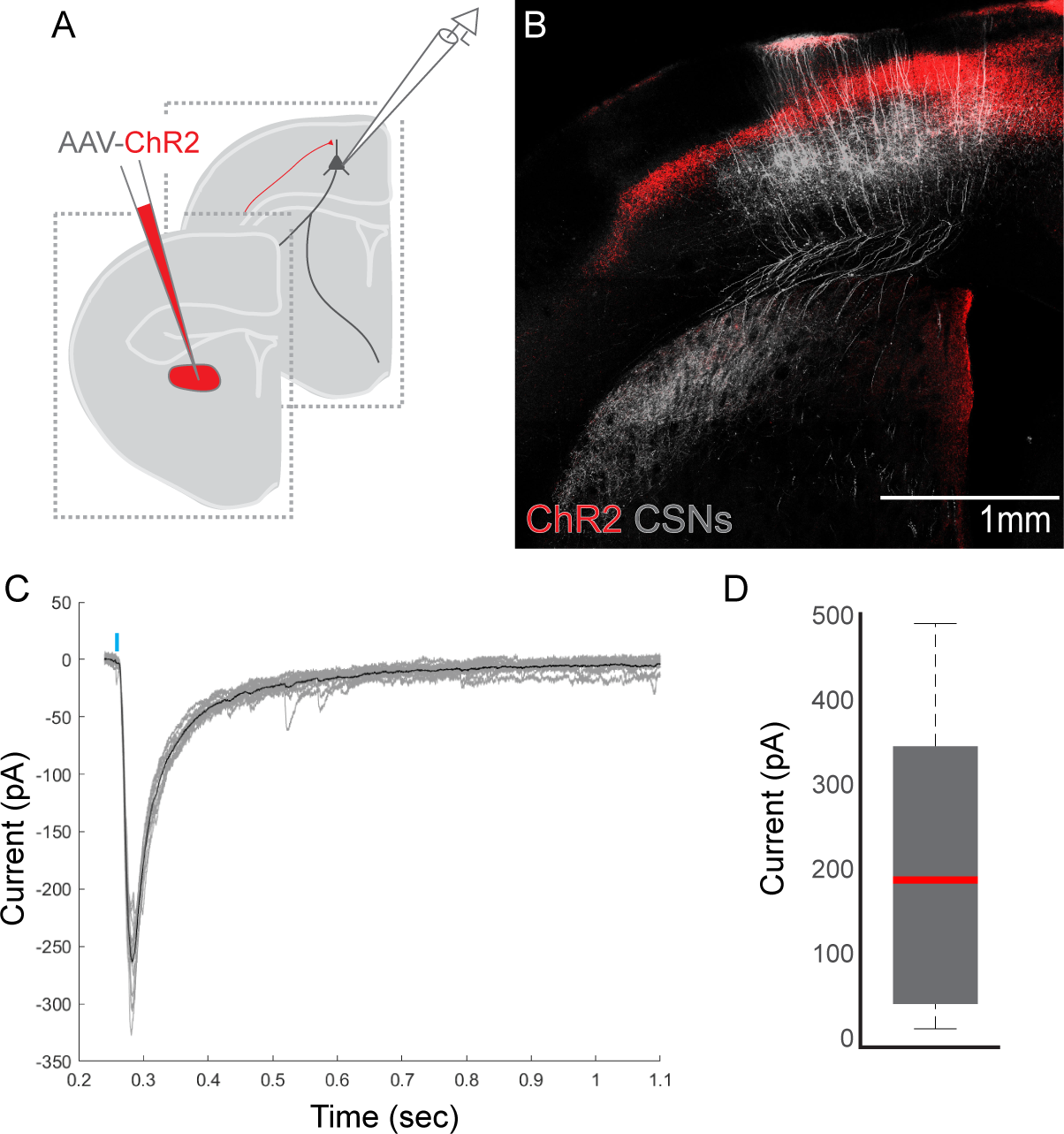
**A)** Strategy to record from CSNs and stimulate thalamocortical projections. **B)** Wide view photomicrograph showing CNs (grey) and thalamocortical axons (red). **C)** Example single trial EPSCs (grey) evoked through ChR2 stimulation (blue bar). The average ChR2-evoked response is overlaid in black. **D)** Boxplot of the ChR2-evoked responses of all CSNs targeted for whole cell recording. The red line indicated the median. The bottom and top edges of the box indicate the 25^th^ and 75^th^ percentiles, respectively. The whiskers extend to the most extreme data points. There were no outliers.

## REFERENCES

1. Arber, S., and Costa, R.M. (2018). Connecting neuronal circuits for movement. Science 360, 1403–1404. 10.1126/science.aat5994.

2. Lemon, R.N. (2008). Descending pathways in motor control. Annu Rev Neurosci 31, 195–218. 10.1146/annurev.neuro.31.060407.125547.

3. Jessell, T.M. (2000). Neuronal specification in the spinal cord: inductive signals and transcriptional codes. Nat Rev Genet 1, 20–29. 10.1038/35049541.

4. Lu, D.C., Niu, T.Y., and Alaynick, W.A. (2015). Molecular and cellular development of spinal cord locomotor circuitry. Front Mol Neurosci 8. ARTN 25 10.3389/fnmol.2015.00025.

5. Russ, D.E., Cross, R.B.P., Li, L., Koch, S.C., Matson, K.J.E., Yadav, A., Alkaslasi, M.R., Lee, D.I., Le Pichon, C.E., Menon, V., and Levine, A.J. (2021). A harmonized atlas of mouse spinal cord cell types and their spatial organization. Nat Commun 12, 5722. 10.1038/s41467-021-25125-1.

6. Porter, R., and Lemon, R. (1993). Corticospinal function and voluntary movement (Clarendon Press ; Oxford University Press).

7. Nelson, A., Abdelmesih, B., and Costa, R.M. (2021). Corticospinal populations broadcast complex motor signals to coordinated spinal and striatal circuits. Nat Neurosci 24, 1721–1732. 10.1038/s41593-021-00939-w.

8. Wang, X., Liu, Y., Li, X., Zhang, Z., Yang, H., Zhang, Y., Williams, P.R., Alwahab, N.S.A., Kapur, K., Yu, B., et al. (2017). Deconstruction of Corticospinal Circuits for Goal-Directed Motor Skills. Cell 171, 440–455 e414. 10.1016/j.cell.2017.08.014.

9. Martin, J.H. (1996). Differential spinal projections from the forelimb areas of the rostral and caudal subregions of primary motor cortex in the cat. Exp Brain Res 108, 191–205. 10.1007/BF00228094.

10. Morecraft, R.J., Ge, J., Stilwell-Morecraft, K.S., McNeal, D.W., Pizzimenti, M.A., and Darling, W.G. (2013). Terminal distribution of the corticospinal projection from the hand/arm region of the primary motor cortex to the cervical enlargement in rhesus monkey. J Comp Neurol 521, 4205–4235. 10.1002/cne.23410.

11. Ueno, M., Nakamura, Y., Li, J., Gu, Z., Niehaus, J., Maezawa, M., Crone, S.A., Goulding, M., Baccei, M.L., and Yoshida, Y. (2018). Corticospinal Circuits from the Sensory and Motor Cortices Differentially Regulate Skilled Movements through Distinct Spinal Interneurons. Cell Rep 23, 1286–1300 e1287. 10.1016/j.celrep.2018.03.137.

12. Levine, A.J., Lewallen, K.A., and Pfaff, S.L. (2012). Spatial organization of cortical and spinal neurons controlling motor behavior. Curr Opin Neurobiol 22, 812–821. 10.1016/j.conb.2012.07.002.

13. Neafsey, E.J., and Sievert, C. (1982). A second forelimb motor area exists in rat frontal cortex. Brain Res 232, 151–156. 10.1016/0006-8993(82)90617-5.

14. Morandell, K., and Huber, D. (2017). The role of forelimb motor cortex areas in goal directed action in mice. Sci Rep 7, 15759. 10.1038/s41598-017-15835-2.

15. Liang, F., Rouiller, E.M., and Wiesendanger, M. (1993). Modulation of sustained electromyographic activity by single intracortical microstimuli: comparison of two forelimb motor cortical areas of the rat. Somatosens Mot Res 10, 51–61. 10.3109/08990229309028823.

16. Kita, T., and Kita, H. (2012). The subthalamic nucleus is one of multiple innervation sites for long-range corticofugal axons: a single-axon tracing study in the rat. J Neurosci 32, 5990–5999. 10.1523/JNEUROSCI.5717-11.2012.

17. Rouiller, E.M., Moret, V., and Liang, F. (1993). Comparison of the connectional properties of the two forelimb areas of the rat sensorimotor cortex: support for the presence of a premotor or supplementary motor cortical area. Somatosens Mot Res 10, 269–289. 10.3109/08990229309028837.

18. Wang, Z., Romanski, A., Mehra, V., Wang, Y., Brannigan, M., Campbell, B.C., Petsko, G.A., Tsoulfas, P., and Blackmore, M.G. (2022). Brain-wide analysis of the supraspinal connectome reveals anatomical correlates to functional recovery after spinal injury. Elife 11. 10.7554/eLife.76254.

19. Tennant, K.A., Adkins, D.L., Donlan, N.A., Asay, A.L., Thomas, N., Kleim, J.A., and Jones, T.A. (2011). The organization of the forelimb representation of the C57BL/6 mouse motor cortex as defined by intracortical microstimulation and cytoarchitecture. Cereb Cortex 21, 865–876. 10.1093/cercor/bhq159.

20. Watson, C., Paxinos, G., Kayalioglu, G., and Heise, C. (2009). Atlas of the Mouse Spinal Cord. Spinal Cord: A Christopher and Dana Reeve Foundation Text and Atlas, 308–379. Doi 10.1016/B978-0-12-374247-6.50020-1.

21. Botta, P., Fushiki, A., Vicente, A.M., Hammond, L.A., Mosberger, A.C., Gerfen, C.R., Peterka, D., and Costa, R.M. (2020). An Amygdala Circuit Mediates Experience-Dependent Momentary Arrests during Exploration. Cell. 10.1016/j.cell.2020.09.023.

22. Zingg, B., Chou, X.L., Zhang, Z.G., Mesik, L., Liang, F., Tao, H.W., and Zhang, L.I. (2017). AAV-Mediated Anterograde Transsynaptic Tagging: Mapping Corticocollicular Input-Defined Neural Pathways for Defense Behaviors. Neuron 93, 33–47. 10.1016/j.neuron.2016.11.045.

23. Zingg, B., Peng, B., Huang, J., Tao, H.W., and Zhang, L.I. (2020). Synaptic Specificity and Application of Anterograde Transsynaptic AAV for Probing Neural Circuitry. J Neurosci 40, 3250–3267. 10.1523/JNEUROSCI.2158-19.2020.

24. Fiederling, F., Hammond, L.A., Ng, D., Mason, C., and Dodd, J. (2021). SpineRacks and SpinalJ for efficient analysis of neurons in a 3D reference atlas of the mouse spinal cord. STAR Protoc 2, 100897. 10.1016/j.xpro.2021.100897.

25. Rexed, B. (1954). A cytoarchitectonic atlas of the spinal cord in the cat. J Comp Neurol 100, 297–379. 10.1002/cne.901000205.

26. Bikoff, J.B., Gabitto, M.I., Rivard, A.F., Drobac, E., Machado, T.A., Miri, A., Brenner-Morton, S., Famojure, E., Diaz, C., Alvarez, F.J., et al. (2016). Spinal Inhibitory Interneuron Diversity Delineates Variant Motor Microcircuits. Cell 165, 207–219. 10.1016/j.cell.2016.01.027.

27. Gabitto, M.I., Pakman, A., Bikoff, J.B., Abbott, L.F., Jessell, T.M., and Paninski, L. (2016). Bayesian Sparse Regression Analysis Documents the Diversity of Spinal Inhibitory Interneurons. Cell 165, 220–233. 10.1016/j.cell.2016.01.026.

28. Zholudeva, L.V., and Lane, M.A. (2023). Spinal interneurons : plasticity after spinal cord injury (Academic Press, an imprint of Elsevier).

29. Mo, A., Mukamel, E.A., Davis, F.P., Luo, C., Henry, G.L., Picard, S., Urich, M.A., Nery, J.R., Sejnowski, T.J., Lister, R., et al. (2015). Epigenomic Signatures of Neuronal Diversity in the Mammalian Brain. Neuron 86, 1369–1384. 10.1016/j.neuron.2015.05.018.

30. Zeisel, A., Hochgerner, H., Lonnerberg, P., Johnsson, A., Memic, F., van der Zwan, J., Haring, M., Braun, E., Borm, L.E., La Manno, G., et al. (2018). Molecular Architecture of the Mouse Nervous System. Cell 174, 999–1014 e1022. 10.1016/j.cell.2018.06.021.

31. Hayashi, M., Hinckley, C.A., Driscoll, S.P., Moore, N.J., Levine, A.J., Hilde, K.L., Sharma, K., and Pfaff, S.L. (2018). Graded Arrays of Spinal and Supraspinal V2a Interneuron Subtypes Underlie Forelimb and Hindlimb Motor Control. Neuron 97, 869–884 e865. 10.1016/j.neuron.2018.01.023.

32. Haring, M., Zeisel, A., Hochgerner, H., Rinwa, P., Jakobsson, J.E.T., Lonnerberg, P., La Manno, G., Sharma, N., Borgius, L., Kiehn, O., et al. (2018). Neuronal atlas of the dorsal horn defines its architecture and links sensory input to transcriptional cell types. Nat Neurosci 21, 869–880. 10.1038/s41593-018-0141-1.

33. Rosenberg, A.B., Roco, C.M., Muscat, R.A., Kuchina, A., Sample, P., Yao, Z., Graybuck, L.T., Peeler, D.J., Mukherjee, S., Chen, W., et al. (2018). Single-cell profiling of the developing mouse brain and spinal cord with split-pool barcoding. Science 360, 176–182. 10.1126/science.aam8999.

34. Sathyamurthy, A., Johnson, K.R., Matson, K.J.E., Dobrott, C.I., Li, L., Ryba, A.R., Bergman, T.B., Kelly, M.C., Kelley, M.W., and Levine, A.J. (2018). Massively Parallel Single Nucleus Transcriptional Profiling Defines Spinal Cord Neurons and Their Activity during Behavior. Cell Rep 22, 2216–2225. 10.1016/j.celrep.2018.02.003.

35. Tashima, R., Koga, K., Yoshikawa, Y., Sekine, M., Watanabe, M., Tozaki-Saitoh, H., Furue, H., Yasaka, T., and Tsuda, M. (2021). A subset of spinal dorsal horn interneurons crucial for gating touch-evoked pain-like behavior. Proc Natl Acad Sci U S A 118. 10.1073/pnas.2021220118.

36. Koch, S.C., Del Barrio, M.G., Dalet, A., Gatto, G., Gunther, T., Zhang, J., Seidler, B., Saur, D., Schule, R., and Goulding, M. (2017). RORbeta Spinal Interneurons Gate Sensory Transmission during Locomotion to Secure a Fluid Walking Gait. Neuron 96, 1419–1431 e1415. 10.1016/j.neuron.2017.11.011.

37. Fink, A.J.P., Croce, K.R., Huang, Z.J., Abbott, L.F., Jessell, T.M., and Azim, E. (2014). Presynaptic inhibition of spinal sensory feedback ensures smooth movement. Nature 509, 43-+. 10.1038/nature13276.

38. Ramón y Cajal, S. (1909). Histologie du système nerveux de l’homme & des vertébrés, Ed. française rev. & mise à jour par l’auteur, tr. de l’espagnol par L. Azoulay. Edition (Maloine).

39. Reardon, T.R., Murray, A.J., Turi, G.F., Wirblich, C., Croce, K.R., Schnell, M.J., Jessell, T.M., and Losonczy, A. (2016). Rabies Virus CVS-N2c(Delta G) Strain Enhances Retrograde Synaptic Transfer and Neuronal Viability. Neuron 89, 711–724. 10.1016/j.neuron.2016.01.004.

40. Hira, R., Ohkubo, F., Tanaka, Y.R., Masamizu, Y., Augustine, G.J., Kasai, H., and Matsuzaki, M. (2013). optogenetic tracing of functional corticocortical connections between motor forelimb areas. Front Neural Circuit 7. ARTN 55 10.3389/fncir.2013.00055.

41. Resta, F., Montagni, E., de Vito, G., Scaglione, A., Allegra Mascaro, A.L., and Pavone, F.S. (2022). Large-scale all-optical dissection of motor cortex connectivity shows a segregated organization of mouse forelimb representations. Cell Rep 41, 111627. 10.1016/j.celrep.2022.111627.

42. Network, B.I.C.C. (2021). A multimodal cell census and atlas of the mammalian primary motor cortex. Nature 598, 86–102. 10.1038/s41586-021-03950-0.

43. Wolff, S.B., and Olveczky, B.P. (2018). The promise and perils of causal circuit manipulations. Curr Opin Neurobiol 49, 84–94. 10.1016/j.conb.2018.01.004.

44. Tomatsu, S., Kim, G., Kubota, S., and Seki, K. (2023). Presynaptic gating of monkey proprioceptive signals for proper motor action. Nat Commun 14, 6537. 10.1038/s41467-023-42077-w.

45. Crapse, T.B., and Sommer, M.A. (2008). Corollary discharge across the animal kingdom. Nature Reviews Neuroscience 9, 587–600. 10.1038/nrn2457.

46. Lundberg, A., and Voorhoeve, P. (1962). Effects from the pyramidal tract on spinal reflex arcs. Acta Physiol Scand 56, 201–219. 10.1111/j.1748-1716.1962.tb02498.x.

47. Warriner, C.L., Fageiry, S., Saxena, S., Costa, R.M., and Miri, A. (2022). Motor cortical influence relies on task-specific activity covariation. Cell Rep 40, 111427. 10.1016/j.celrep.2022.111427.

48. Drew, T., Jiang, W., and Widajewicz, W. (2002). Contributions of the motor cortex to the control of the hindlimbs during locomotion in the cat. Brain Res Brain Res Rev 40, 178–191. 10.1016/s0165-0173(02)00200-x.

49. Kamiyama, T., Kameda, H., Murabe, N., Fukuda, S., Yoshioka, N., Mizukami, H., Ozawa, K., and Sakurai, M. (2015). Corticospinal tract development and spinal cord innervation differ between cervical and lumbar targets. J Neurosci 35, 1181–1191. 10.1523/JNEUROSCI.2842-13.2015.

50. Moreno-Lopez, Y., Bichara, C., Delbecq, G., Isope, P., and Cordero-Erausquin, M. (2021). The corticospinal tract primarily modulates sensory inputs in the mouse lumbar cord. Elife 10. 10.7554/eLife.65304.

51. Peters, A.J., Lee, J., Hedrick, N.G., O’Neil, K., and Komiyama, T. (2017). Reorganization of corticospinal output during motor learning. Nat Neurosci 20, 1133–1141. 10.1038/nn.4596.

52. Currie, S.P., Ammer, J.J., Premchand, B., Dacre, J., Wu, Y.F., Eleftheriou, C., Colligan, M., Clarke, T., Mitchell, L., Faisal, A.A., et al. (2022). Movement-specific signaling is differentially distributed across motor cortex layer 5 projection neuron classes. Cell Reports 39. ARTN 110801 10.1016/j.celrep.2022.110801.

53. Barrett, J.M., Martin, M.E., and Shepherd, G.M.G. (2022). Manipulation-specific cortical activity as mice handle food. Curr Biol 32, 4842–4853 e4846. 10.1016/j.cub.2022.09.045.

